# A Discrete Formulation of Two Alternative Forced Choice Decision Dynamics Derived from a Double-Well Quantum Landscape

**DOI:** 10.1101/2021.09.23.461524

**Authors:** Morgan Rosendahl, Jonathan Cohen

## Abstract

Tools from quantum theory have been effectively leveraged in modeling otherwise poorly understood effects in decision-making such as apparent fallacies in probability judgments and context effects. This approach has described the dynamics of two alternative forced choice (2AFC) decisions in terms of the path of a single quantum particle evolving in a single potential well. Here, we present a variant on that approach, which we name the Multi-Particle and Multi-Well (MPMW) quantum cognitive framework, in which decisions among N alternatives are treated by the sum of positional measurements of many independent quantum particles representing stimulus information, acted on by an N-well landscape that defines the decision alternatives. In this article, we apply the MPMW model to the simplest and most common case of N-alternative decision making, 2AFC dynamics. This application calls for a multi-particle double-well implementation, which allows us to construct a simple, analytically tractable discrete drift diffusion model (DDM), in the form of a Markov chain, wherein the parameters of the attractor wells reflect bottom-up (automatic) and top-down (control-dependent) influences on the integration of external information. We first analyze this Markov chain in its simplest form, as a single integrator with a generative process arising from a static quantum landscape and fixed thresholds, and then consider the case of multi-integrator processing under the same conditions. Within this system, stochasticity arises directly from the double-well quantum attractor landscape as a function of the dimensions of its wells, rather than as an external parameter requiring independent fitting. The simplicity of the Markov chain component of this model allows for easy analytical computation of closed forms for response time distributions and response probabilities that match qualitative properties of the accuracies and reaction times of humans performing 2AFC tasks. The MPMW framework produces response time distributions following inverse gaussian curves familiar from previous DDM models and empirical data, including the common observation that mean response times are faster for incorrect than for correct responses. The work presented in this paper serves as a proof of concept, based on which the MPMW framework can be extended to address more complex decision-making processes, (e.g., N-alternative forced choice, dynamic control allocation, and nesting quantum landscapes to allow for modeling at both the task and stimulus levels of processing) that we discuss as future directions.

## 1 Introduction and Background

### 1.1 Description of the 2AFC Paradigm

Forced-choice decision making under uncertainty and time constraints has been studied and modeled extensively throughout decades of work in psychology, neuroscience, economics, and other disciplines (see [4],[28],[24],[16]). This has been most commonly studied in the context of 2AFC decision making. Although this is a simplification of the sorts of choices agents often face, it has proven useful as a paradigm that is tractable to formal analysis. Many formal models have emerged to address 2AFC decision making and one, the continuous time drift diffusion model (DDM) ([11]; [18]; [25]), has emerged as the gold standard among them. The DDM formalizes the 2AFC problem as choosing the stronger of two noisy signals when signal strength is unknown, either when forced to choose at a time determined by an instruction (e.g., from the experimenter; a process termed the interrogation paradigm) or when sufficient evidence for one of the alternatives has accumulated (a process termed the free response paradigm). Information about the signals is accumulated by integrators (e.g., neural populations) differentially responsive to the competing alternatives. The dynamics of the decision-making process, defined as the process by which information is presented to and integrated by an agent, are described by reducing two integrators responsive to evidence for each alternative separately to a single integrator that represents the difference in the accumulated evidence.

The power of the DDM is that, despite its relative simplicity, it captures important qualities of performance as well as neural signatures of the decision process in 2AFC tasks in both humans and other species, including right skewing response time distributions and asymmetry in expected response times between correct and incorrect responses. In the next section, we briefly review the continuous DDM and a founding quantum walk model analogous to it, then use the two to frame the model presented in the remainder of this article.

### 1.2 The DDM as an Optimal Solver of 2AFC Tasks

The continuous DDM [18] describes the stochastic decision-making process by a state variable, *x*(*t*), as the sum of a series of independent identically distributed (IID) Gaussian variables. The resultant change in state is given by

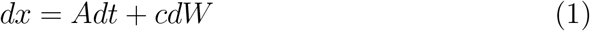

where *A* is the mean drift rate, representing signal strength, and the second term is Gaussian distributed noise. Note that this model, like many others, includes noise as a free parameter. This represents a continuous implementation of the sequential probability ratio test ([32]; see [18]), and therefore an optimal procedure for solving 2AFC problems: for a specified accuracy, it guarantees the fastest decision time and, conversely, for fixed decision time it guarantees the highest accuracy. That is, this procedure defines a pareto relationship between the time required to reach a decision with the accuracy of the decision reached, known commonly as “the speed-accuracy tradeoff.”

Other models for 2AFC decision-making have been proposed that introduce additional factors and corresponding parameters. Such models include the Ornstein-Uhlenbeck integration process ([4], the Leaky Competing Accumulator (LCA) model that relies on mutual inhibition for differencing ([28]) and ones that implement decisions through pooled inhibition([33]). However, Bogacz et al ([1]) have shown that choosing the parameters that optimize 2AFC performance for the speed-accuracy tradeoff reduces these models to the DDM, reaffirming its value as a foundational model of 2AFC decision dynamics.

As noted above, the DDM defines a family of speed-accuracy values, that vary as a function of the threshold at which the integration process is halted and a decision is reached. While these are optimal with regard to one factor (time or accuracy) given the other, in a given setting an agent must decide which to prioritize and by how much. This can be formalized as another optimization problem, using reward rate (i.e., accuracy per unit time) maximization as the objective function. This amounts to determining the threshold that yields the maximum reward in minimal time within the constraints of specific task conditions. This is a natural objective function, that has been used to explain the pattern of speed-accuracy tradeoffs observed in studies of human behavioral performance ([1]), and comports with the instruction commonly provided in such studies to “respond as quickly and accurately as possible” on each trial.

### 1.3 The Quantum Walk 2AFC

The application of quantum mechanical formalisms to human decision-making processes in the form of a random walk was introduced by Busemeyer and colleagues ([7]) as one component of a broader approach applying quantum principles to the study of cognition. This work was motivated by the observation that cognitive and behavioral phenomena share several important properties with quantum systems, and that quantum tools might be suited to explaining some ways in which cognitive processes appear to violate classical laws of probability. One shared property is that both are stochastic. Traditional models of cognition treat stochasticity as a form of noise (e.g,. the influence of factors uncorrelated with the process of interest), the effects of which are a free parameter that must be estimated. In contrast, stochasticity emerges as a fundamental property of quantum systems. Another property shared by cognitive and quantum systems is non-commutativity of measurements. That is, taking a measurement of the system perturbs it, affecting future measurements of the system. In a cognitive system, this process describes order effects: the fact that the order in which measurements are made (e.g., in which an agent is asked questions or to perform a sequence of tasks) effects performance (e.g., the answers they give or the way in which they perform the tasks). In a physical system, this occurs because measuring a system’s state collapses it from one distributed across all possible values to one temporarily described by a single value. Quantum approaches to cognition seek to exploit the analytic approaches developed for use in quantum mechanics to unify these quantum-like cognitive effects with other cognitive phenomena that appear to violate classical laws of probability in a singular model.

One example of a successful application of quantum mathematics to cognitive phenomena takes advantage of the fact that, in quantum systems, a form of interference effect arises from the treatment of statistics in a quantized and vector-based form, in contrast to the classical treatment of statistics as scalar-based. This difference can account for a number of phenomena, such as conjunction-disjunction effects, in which human estimates of probability violate the law of total probability,([15]). Another example from the work of Busemeyer and colleagues ([5], [15], showed that the effects of categorization on decisions, in which differences in choice probabilities arise from making a relevant categorization before versus after the choice, are better described by quantum interference effects than classical probability.

Quantum cognitive models propose that, like physical quantum mechanical systems, decision processes can be described as occupying a state space spanned by a set of “basis states”. The quantum random walk 2AFC model ([7]) defined the basis states as corresponding to levels of preference for each of two alternatives, and their corresponding outcome. Confidence levels are defined as regions along the width of a sloped, one-dimensional attractor well that occupies the x-axis, with total confidence in either alternative at either boundary of the well. An attractor well describes an area to which particles are attracted and within which they may be bound, as by gravity. A sloped attractor well is one for which attraction increases from one side of the well to the other, driving a particle from its starting point to the point at which the potential is strongest (i.e., at which the well is “deepest”, like a ball rolling down a ramp) at one of its edges. The decision process begins when a particle is prepared in a neutral basis state, close to the center of the well and allowed to evolve in accordance with the Time-Dependent Schrodinger Equation (TDSE). Much like the classical continuous DDM, the particle’s state diffuses from its initial state over time while also drifting toward the well boundary with the strongest potential value (determined by task information, such as signal strength). Its state at any time is described by a probability distribution function (PDF) that is distributed across all possible basis states, spanning the entire width of the well.

The classical DDM models the free response paradigm as the agent repeatedly sampling the stochastic stimulus to update the state of the decision process, making a decision when it has reached one of the decision thresholds. Busemeyer et al ([7]) model the agent’s updates of their decision process by repeatedly measuring the position of a quantum particle within the well, where its position determines the agent’s confidence level. Prior to measurement, as stated above, quantum particles exist in a state distributed across all points within the well, described by their PDF. Upon measurement, the state collapses to the point at which the particle is measured. The confidence level within which the particle is measured defines the state of the agent’s decision process at that time, and the particle continues to evolve under the TDSE from that state. An agent sets decision boundaries at certain confidence levels and, if the confidence level that the particle falls within is greater than that at which the decision boundary has been set, the agent determines that they have reached a decision and reports it. The probability that an agent makes a decision for one or the other alternative at any time is determined by finding the component of the PDF that occupies the region between the decision boundaries and the well boundaries, which gives the probability that the agent is at least as certain of their choice as they need to be to make a decision.

An agent may set their decision boundaries at different confidence levels to optimize reward rate, just as in the classical DDM. For example, an agent that favors accuracy over speed may set their boundaries at 98% confidence, while an agent more concerned with speed than accuracy may set them at 80%. As yet, however, threshold optimization has not been addressed within the quantum walk 2AFC model.

Although this formulation of the quantum walk is similar to the classical DDM, it makes some novel predictions. Because the motion of the system is that of a quantum particle, over time, different components of the particle’s state will reach the well boundaries and reflect back into the well, “sloshing” like a classical wave, and producing interference in the state’s PDF. Thus, under the interrogation paradigm, the state of this system will never permanently settle on one decision or another. Instead, it will continue to oscillate between the boundaries in accordance with the TDSE; an agent interrogated repeatedly over a long period of time will gradually and repeatedly change their mind in accordance with the interference and oscillations of the quantum state. In experimental work designed to test this theory, the Busemeyer group found evidence in favor of such oscillations, but also the need to combine their quantum walk with a markov approach in order to better capture the effect. To do so, they have recently developed an extension to their previous quantum random walk model, an open system model of preference-based decision-making, combining quantum and classical approaches [13] to explain damped oscillations in preference formation.

Following the framework of the quantum random walk 2AFC, a number of quantum cognitive models, such as those presented in Quantum Models of Cognition and Decision ([15]), are built upon the repeated measurements of a single particle within a quantum potential, made by asking the agent to report their mental state, thus perturbing the system and affecting future measurements. These models have been used to successfully explain order effects ([27]; [34]; [35]) in value-based judgment and errors in probability judgment ([17]).

### 1.4 The Multi-Particle Multi-Well (MPMW) Model

In the remainder of this article, we will describe the MPMW, which was first introduced in [22]. The MPMW framework was constructed for the purpose of modeling perceptual forced choice decision dynamics, utilizing both quantum and classical components. The MPMW differs from other quantum models in several of its formalisms. First, the MPMW takes a multi-well approach, assigning a single well to each stimulus-response alternative, that makes extending the MPMW to greater than two alternatives relatively easy. Additionally, it treats multiple, independent, sequentially processed particles, which allows the decision dynamics to be analyzed using a simple Markov chain (or set of markov chains, in the case of greater than two alternatives). The MPMW also makes use of a broader range of tools from quantum mechanics to capture a richer variety of phenomena. These tools include the use of eigenenergies to model arousal and particles’ access to the “classically forbidden region” (the region outside of wells, a phenomenon unique to quantum mechanics) as fundamental to the model. Furthermore, because the MPMW operates using sequentially sampled particles in discrete time, it places an upper bound on the amount of information that can be processed by an agent at any given time (manifesting as bounds on drift rate and variance). That is, unlike in the case of continuous time, continuous state models, the MPMW does not allow an infinite amount of evidence to be integrated in infinitesimal time. This bound causes the MPMW to make predictions that differ from the DDM in certain cases (e.g., the case of a small number of integrators and high conflict among alternatives). Finally, the MPMW is easily extensible to treat multiple parallel integration processes. This feature allows the MPMW to reproduce predictions that are similar to those of the DDM by its convergence to a DDM in the limit of a large number of integrators.

In the quantum cognitive models summarized above, the basis states are defined as sections along the width of the attractor well. However, in physical quantum systems, it is more common to define the basis states by the discrete set of particle energies that may occupy the well, as determined by its dimensions and boundary conditions of the Schrodinger Equation. These characteristic energies, referred to as eigenenergies throughout the remainder of the paper, define the range of energies at which emitted particles have nonzero probability of being measured within the attractor wells and are found by solving the Time-Independent Schrodinger Equation (TISE) for the system. One drawback of the quantum random walk as derived by Busemeyer et. al ([7]) is that it does not account for or incorporate the eigenenergies, thus ignoring an important feature of quantum systems. In the MPMW framework, eigenenergies can be thought of as representing the level of arousal of the agent, that determines how efficiently it can integrate task-relevant information. A thorough exploration of the effects of well parameters on eigenenergies and eigenenergies’ consequences for information intregration is not within the scope of this article, but is covered in related work (Rosendahl & Cohen, Preprint). Briefly, we note that lower eigenenergies, representing a lower arousal (”calmer”) state, correspond to more efficient information integration, and thus improved efficacy of applied control. Importantly, increasing stimulus salience (well width) or control (well depth) increases the number of available eigenenergies, thus making information integration more robust to perturbations in arousal energy, and cause the eigenenergies to be associated with higher integration efficiency, thus producing faster response times with a lower error rate.

Another important feature of quantum systems not addressed by the quantum walk 2AFC model is the ability of a quantum particle to be found outside the well in the “classically forbidden region”, so named because it is not accessible in classical systems without a change in energy. In physical quantum systems, it is possible for a particle to enter the classically forbidden region without a change in energy by a process known as quantum tunneling. However, in the Busemeyer et al ([7]) model, it is assumed that the particle never leaves the well, using renormalization to permanently bind its probability distribution function within the well walls. In the MPMW model, particles may access the classically forbidden region and particles that are found there are treated as information that is effectively lost to the system, as it falls outside of the landscape components relevant to the decision-making process. The probability that a particle is found in the classically forbidden region is an emergent property of the solutions to the TISE, and is therefore bound inextricably to the dimensions of the attractor wells in a landscape. Thus, by the use of the classically forbidden region, the MPMW features a form of stochasticity (i.e., “noise”) that is emergent of context and does not need to be fitted as a free parameter of the model. Particles measured in the classically forbidden region are treated as “dropped bits”, and do not advance the decision variable toward any boundary; they serve a purpose very similar to “leak” in classical models.

In the sections that follow, we present an implementation of the MPMW 2AFC model that parallels the quantum random walk ([7]) model. The MPMW framework, applied to model perceptual forced choice problems, includes as many wells as competing stimuli, incorporates the effects of their associated eigenenergies, reintroduces the accessibility of the classically forbidden region, and models the information integration process by measuring those effects on multiple independent particles. The MPMW’s change in approach provides three important advantages, by allowing the model to: a) be extended beyond 2AFC tasks to ones with any number of alternative; b) provide a natural interpretation of automaticity or stimulus salience and the effects of control in terms, respectively, of the width and depth of individual attractor wells, and c) address effects of distraction or fluctuations in arousal in terms of fluctuations in eigenenergy that increase the rate at which particles occupy the classically forbidden region. Here, we implement a version of the MPMW that addresses decision making in 2AFC tasks, that allows us to compare it directly to the classical DDM and quantum random walk approach. In other work (currently unpublished) we have begun to explore the unique characteristics listed above.

In its simplest form, that we explore in this article, the MPMW models the 2AFC task as a generative process emitting particles within a constant range of energies into a static double-well landscape. We show that this can be reduced to a classical discrete-time Markov chain that is analytically tractable and yields similar results to the classical DDM while incorporating important features of quantum models and making some predictions that differentiate it from the DDM at low numbers of integrators and at optimal thresholds. Following its description, we analyze the predictions the MPMW model makes when applied to the interrogation and free response paradigms, as both a single integrator system, and a multiple integrator system in which the outcome of the decision process reflects the sum of a population of integrators. We will also derive optimal thresholds to maximize reward rate under different reward schema as a function of drift rate and number of integrators.

## 2 Introduction to the Double-Well Implementation of the MPMW Model for 2AFC Tasks

To motivate the 2AFC implementation of the MPMW framework, consider a dot motion task: an agent is presented visually with a field of dots, the majority of which are moving randomly, but a subset of which are moving coherently left or right. The agent is tasked with classifying the prevailing direction of motion as “left” or “right” and making a corresponding response (e.g., press ‘F’ on a keyboard for left, and ‘J’ for right; [2]; [23]). Correct answers are rewarded while incorrect answers are penalized, each by known amounts that may not be equal. We model the incident stimulus at each point in time as a quantum particle emitted into a landscape with two square attractor wells representing each of the two response alternatives. The width of each well is determined by the salience of its associated stimulus component (e.g., if the prevailing motion is left, the left-associated well will be wider than the right-associated well), and the depth of the wells is determined by the degree of control (e.g., attention) allocated to attending that stimulus component (in this case, we would expect the depths to be equal, as there is no reason for the agent to prefer attending left-moving over right-moving dots) and binding that stimulus to its assigned response (e.g., if left, press ‘F’ on the keyboard. if right, press ‘J’).

As in all quantum systems, prior to its measurement, the state of each particle is distributed across the two wells and the classically forbidden region. This models the distribution of the stimulus representation as it is initially presented to an integrator, containing information about all of the dots, the subtleties of their motion, and irrelevant stimuli in the agent’s field of view in that moment. In order to make the decision, this complex representation must be reduced to a single category (left or right). The integrator performs this categorization of the stimulus as it appears at each point in time, and each categorization (left, left, right, left) contributes to the accumulated evidence, advancing the decision variable toward one or the other decision threshold. This is modeled as a measurement of each particle’s position as they are emitted into the system, one at a time. Performing the measurement collapses each particle’s state to a single point in one of the attractor wells or the classically forbidden region. When a particle’s collapsed state is within the well associated with the response “left” (or “right”), the agent will accumulate a single “bit” of evidence for this alternative, driving the decision variable toward a corresponding decision threshold. If a particle’s collapsed state is within the classically forbidden region, it is considered a “dropped bit” (i.e., information accumulated in a way that is irrelevant to the task) and influences the decision process in the way that leak does in other formulations, slowing the process, but not advancing the decision variable toward either decision threshold.

The probability that a particle is found in each of the three possible states (well 1, well 2, or the classically forbidden region) is found by taking the L2-norms (the continuous analog of a dot product) of the particle’s eigenstates within the boundaries of each well. These states are determined by the parameters of the potential landscape (the widths, depths, and separation of the attractor wells). Modeling the system in this way, stochasticity (i.e., noise) emerges as a fundamental property of the formalism that is coupled directly to stimulus properties and the agent’s familiarity with (automaticity) and control (attention) allocated to the task, thus removing the need introduce noise as a free parameter.

In summary, according to the MPMW framework, the decision process can be thought of as the sequential measurement of particle positions by an array of particle detectors placed throughout the potential landscape. A particle is emitted by the stimulus and its distributed state is determined by the potential landscape: a set of attractor wells shaped by stimulus properties, the agent’s prior experience, and their allocation of control, sitting within a broader landscape (the classically forbidden region). Its state is collapsed when measured (categorized as evidence for one of the alternatives), and then cleared from the system by the measurement process. We model the sequential integration of information (i.e., evidence accumulation) in discrete time by individual particles being emitted, measured, and cleared. The system proceeds in this way, collecting evidence for each alternative, and representing the decision variable as the difference between the two sets of accumulated evidence, until reaching a decision threshold. Treating time as discrete separates this model from many other perceptual decision-making models, which have made the reasonable assumption that time is continuous. Following this assumption, continuous time models were derived by taking discrete time models to the limit of infinitesimal time steps ([8]). While it is reasonable to think of time itself as continuous, a growing body of evidence suggests that perception of attended stimuli occurs in discrete temporal increments. For example, this has motivated the blinking spotlight of attention model, as summarized by VanRullen, Carlson, & Cavanaugh ([30]), that has received considerable support from neural and behavioral data ([31]; [3]; [29]). Furthermore, treating the information accumulation process as discrete places an upper bound on the rate at which information can be accumulated, rather than allowing infinite information to be accumulated in infinitesimal time, thus offering a constrained, more biologically plausbile model of the process.

## 3 Theoretical Analysis and Results

A distinguishing characteristic of the MPMW model is that, rather than sampling a single particle repeatedly (as in [7]), it treats the decision process as a sequential sampling of multiple particles, measuring each only once before it clears the system. Using this sampling paradigm allows us to reduce the MPMW model to a classical Markov chain with analytical expressions for probabilities of each choice and response time distributions.

The qualitative properties of the Markov chain process are very similar to a traditional DDM, which was derived by taking a discrete-time discrete-state Markov chain to its continuous limit ([8]), but with greater ease of analysis and some interference properties similar to those exhibited by the quantum random walk ([7]). Here, we apply the model to a classic 2AFC problem to find the probabilities of success under the interrogation paradigm and derive closed form solutions for reaction times in the free response paradigm. We show that, in the limit of large integration time, which is mathematically equivalent to continuous time as single time steps become much smaller than integration time, the PDF of the decision variable becomes Gaussian and the RT distributions of this model are given by inverse gaussian distributions comparable to those exhibited by the DDM. We carry out an identical analysis to show that the same is true for a multiple integrator system in the limit of large number of integrators. We also analyze key aspects of the model’s behavior that differ from both the traditional DDM and single particle quantum models, thus providing an opportunity to distinguish among these in future empirical studies.

### 3.1 The Single Integrator Model

#### 3.1.1 The State of the System Over Time

In the MPMW 2AFC model, the displacement of the decision variable *X*(*t*) is the sum of IID variables, the probability distributions of which are given by the probability of an incident particle (or bit) being accumulated as evidence for either response 1, response 2, or neither.

Value assignments for each bit take on the value of a step size for a random walk, given by

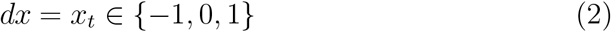

and the agent’s progress from its initial position at time *τ* is given by

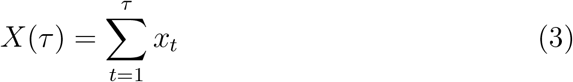

Note that this is a significant departure from the traditional DDM, in which step size is unbounded and decisions may, therefore, be reached instantaneously. In the single-integrator MPMW, not only is step size bounded between −1 and 1, but so is drift rate for a single integrator, enforcing a minimum response time.

The probability of each value of *x_t_* is drawn from a uniformly distributed set of probabilities, which arise from a set of equally probable bound eigenstates admitted by the quantum landscape,*V* (*d*_1_, *d*_2_, *w*_1_, *w*_2_, *w_b_*) (this assumption of equally probable eigenstates is relaxed in later work). Variables denoted *d_i_* give well depths, variables denoted *w_i_* give well widths, and the variable *w_b_* denotes the width of the barrier separating the two wells. We denote the probability that step *x_t_* has a value *a*, given the eigenstate *j* as,

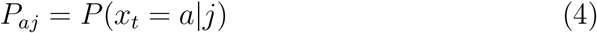

where *j* ∈ [1, 2, 3, *…n*] is the index of the substate.

The expectation of *x_t_*, analogous to the mean drift rate of the standard DDM, is therefore given by

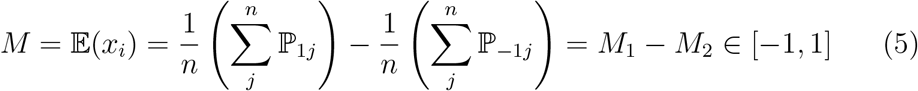

with variance

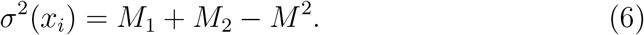

We refer to the terms *M*_1_ and *M*_2_ as the Mean Integration Efficiencies (MIEs) of each well.

The stochastic trajectory of this model arises directly from the competition between target and distractor stimulus-associated attractor wells in the landscape, each of which exerts an attractive force on each particle, laying claim to some part of its pre-measurement probability distribution.

This double-well version of the MPMW is a discrete-time, discrete-state quantum variant of the DDM, that evolves as a Markov chain with a (possibly infinite) single-step transition matrix. This gives the probability of the decision-variable transitioning from state *i* to state *j* in a single step, as

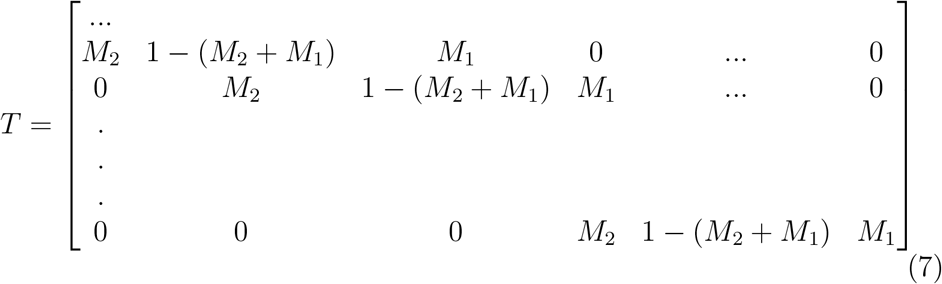

Decision bias can be implemented in one of two mathematically equivalent ways: setting the initial state *X*(0) ≠ 0 or setting unequal boundaries for each alternative. In the quantitative analysis below, we implement the second of these. In principle, drift rates can also vary within and between trials by a combination of the relative widening of a well (assumed to represent stimulus salience or prior experience and the corresponding effects of automaticity), the relative deepening of a well (assumed to represent the effects of control, manifested as the purposive allocation of attention or inhibition of a non-target stimulus-response pairing), or fluctuations in the range of eigenenergies (caused by fluctuations in agent arousal). Although modeling systems exhibiting variable drift rates is outside of the scope of this paper, we will address it in future work.

In the sections that follow, we analyze the two common approaches to 2AFC tasks: the interrogation paradigm and the free response paradigm. Under the interrogation paradigm, the agent is exposed to the stimulus until a time determined by the experimenter, and then forced to respond. Under the free response paradigm, the agent is instructed to respond as quickly and accurately as possible, then exposed to the stimulus until they decide to respond, assumed to occur when the system hits one of two decision thresholds. Under the interrogation paradigm, we are concerned only with accuracy (i.e., probability of a correct choice vs. errors). Under free response, we are interested in both accuracy and response time, as well as the tradeoff between these, and the extent to which it maximizes reward rate. For the MPMW, the interrogation and free response paradigms call for different analyses.

#### 3.1.2 The Interrogation Paradigm

Under the interrogation paradigm, the systemis allowed to evolve freely and its state measured at a time determined by the experimenter,*τ*. As with the standard DDM, the probability of success is given by

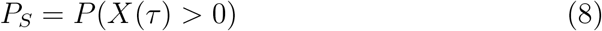

In order to find these values, we need to know the probability that the system is in a state *X_i_* given that it started in state *X*_0_. As noted above, the bounds on the system’s step size, *x_t_*, place equivalent bounds on the state of the system, which may fall into the range

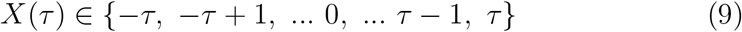

We find the PDF of the decision variable’s state by fixing the size of the transition matrix such that *T* ∈ *R^n×n^*

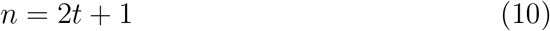

and finding

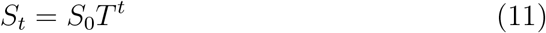

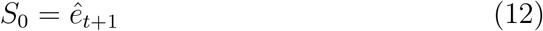

where *ê*_*t*+1_ ∈ *R*^1*×n*^ is the characteristic vector (also called one-hot vectors or standard basis vectors), with all entries zero except the *t* + 1^*st*^ entry, which is 1, and corresponds to starting the system at postion *x* = 0 time *t* = 0. *S_t_* is the probability distribution for all possible states of the system,*X*(*t*).

*T* is a finite tridiagonal matrix with constants on the center diagonal as well as the +1 and −1 diagonals. This type of matrix, known as a Toeplitz tridiagonal, allows for analysis of arbitrary positive integer powers ([12]). In our case, the expressions for the values of the positive integer power matrices can be simplified by algebraic manipulation, arising from the fact that the three constants are determined by two variables, *M*_1_ and *M*_2_. They become,

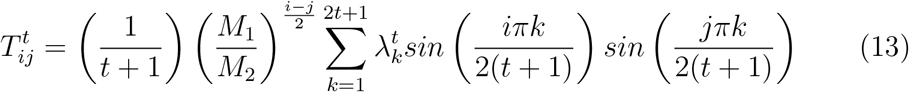

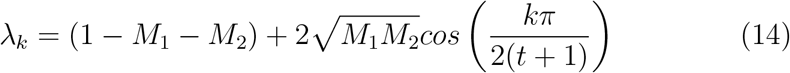

Since the system began at *t* = 0, its probability distribution at time *t* is described by the entries of the *t* + 1^*st*^ column of *T^t^*, given by

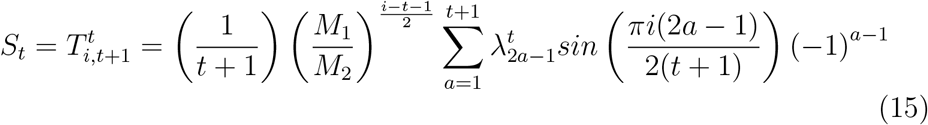

This state, *S_t_*, is a gaussian-like distribution with period 2 interference, arising from the fact that the initial PDF is, effectively a tripartite wave, the components of which interfere with one another. As time progresses, the interference dissipates, as illustrated in figure 1. In section 3.1.4, we will show that, as the interference dissipates, the PDF converges to a gaussian distribution, recovering the functional form of a traditional continuous DDM. This is not a mathematical coincidence, but occurs because taking the PDF of the decision variable in the limit of large time is mathematically equivalent to the process used to derive the continuous DDM ([8]).

**Figure 1:**
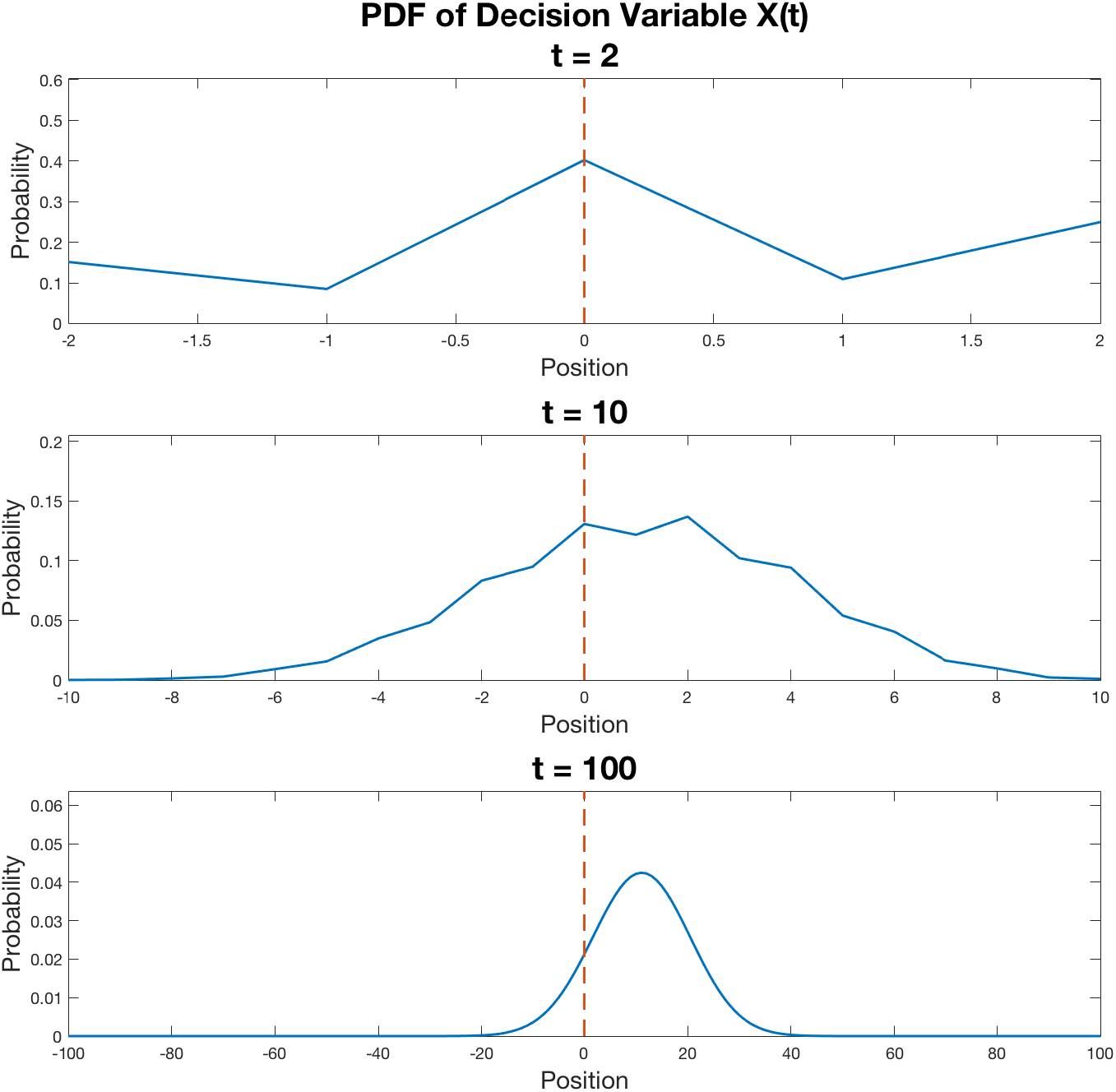
PDF of the decision variable for a system with *M*_1_ = 0.5, *M*_2_ = 0.39 at several interrogation times.

Without loss of generality, we assign the positive direction to be associated with the correct response, and find that the probability of success is given by

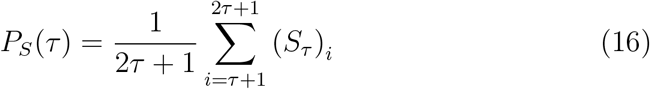

As shown in figure 2, the probability of success as a function of interrogation time is overall increasing (as long as *M*_1_ > *M*_2_), but oscillates with period 2, just as the state of the system does. The amplitude of these oscillations decays over time and is smaller for higher drift rates, as the PDF moves more rapidly toward the decision thresholds.

**Figure 2:**
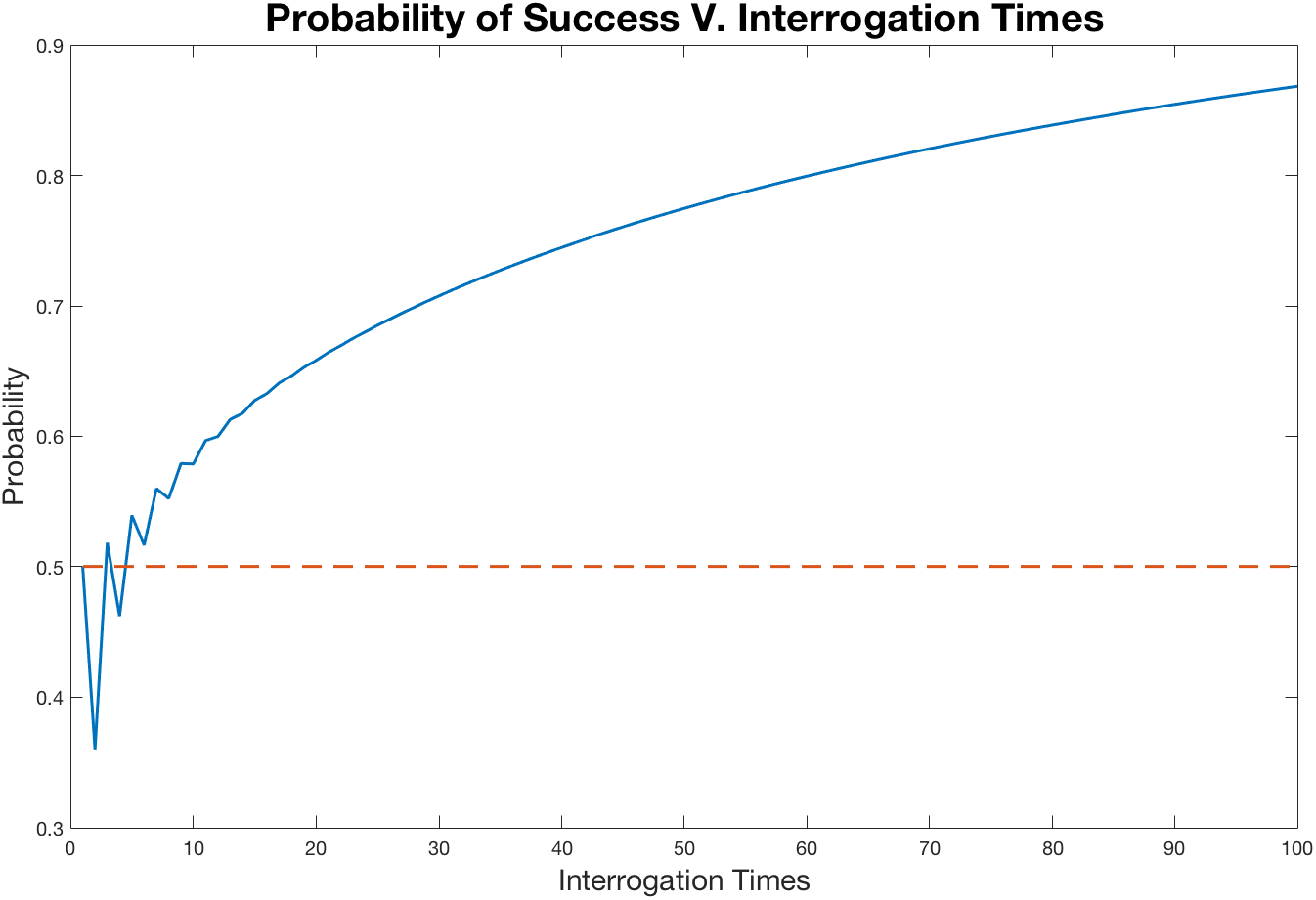
Probability of success upon interrogation for a system with *M*_1_ = 0.5, *M*_2_ = 0.39 across 100 interrogation times.

We perform a numerical analysis of the behavior of the envelope enclosing these oscillations in section 4.1.1.

#### 3.1.3 The Free Response Paradigm

Under the free response paradigm, we set two decision thrsholds at *x* = {−*T_f_, T_s_*}, where *T_s_* is the threshold for making the correct decision (success) and −*T_f_* is the threshold for making the incorrect decision (failure). We do not assume that these thresholds are equal, which corresponds to allowing for the existence of starting point bias in a standard DDM. The system then evolves as a Markov chain with two absorbing states. Its one-step transition probability matrix, giving the probability of transitioning from state *i* to state *j*, is given by

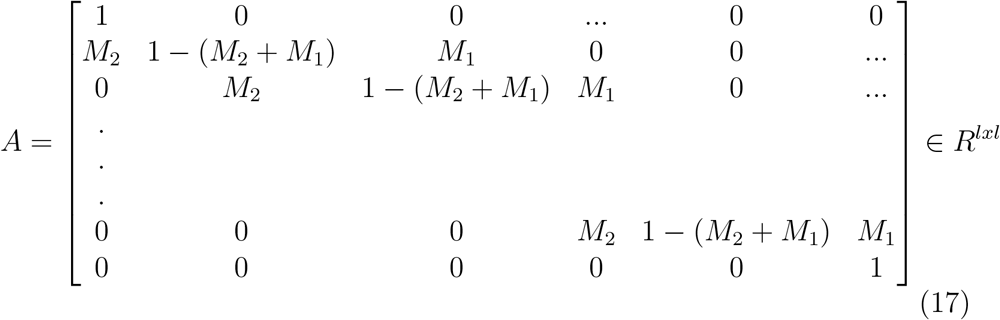

where

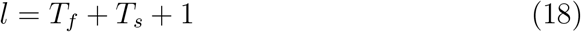

The system may occupy any state in a discrete space given by *x* ∈ *R^l×^*^1^. The state of the system at time *t* is again given by

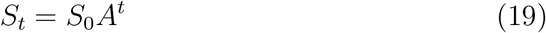

The probabilities of success and failure can be described by setting the decision thresholds as absorbing states and finding the probability of absorption by each from the starting point.

The total probabilities of success and failure are found by solving the recursion relations

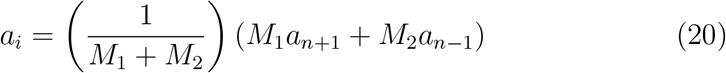

with constraints

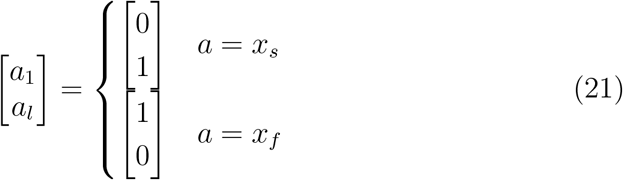

The entries (*x_s_*)_*i*_ give the probability of success, having started from *x*_0_ ∈ [−*T_f_, −T_f_* + 1, *…T_s_* − 1, *T_s_*], and failure is given likewise by (*x_f_*)_*i*_. The boundary conditions state that, if the agent begins exactly at the threshold for the correct decision, their probability of success is 1 and failure is 0, and conversely if it begins at the threshold for an incorrect decision.

The entries of *x_s_* and *x_f_* are exponential and given by

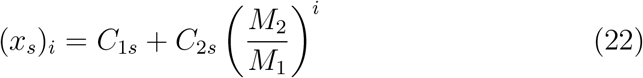

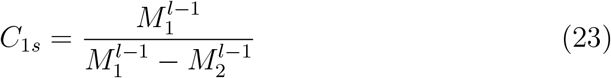

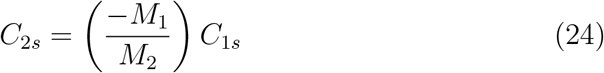

and

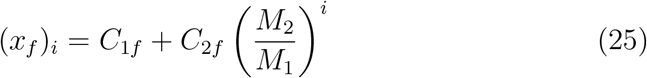

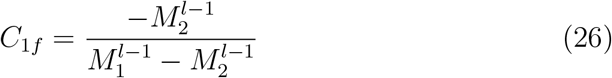

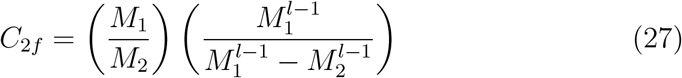

With these values in hand, we can now set the initial position of the agent to *x*_0_ = 0, and the probability of success and failure are given by the 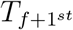 entries of *x_s_* and *x_f_*.

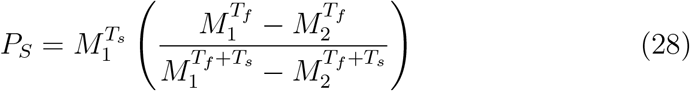

The probability of the agent choosing incorrectly is defined and found similarly as

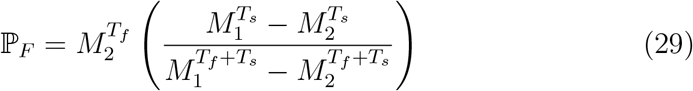

where *M*_1_ ≠ *M*_2_. Where *M*_1_ = *M*_2_, *P_S_* = *P_F_* = 0.5. Success rates across batches of trials follow a binomial distribution, and the probability of success is a sigmoidal function of drift rate, recognizable as the common psychometric curve. The gain of this curve is determined by the absolute value of the thresholds and its bias is determined by the difference in the absolute values of the thresholds. Note that, unlike standard psychometric curves, which are symmetric, the sigmoid curve predicted by the MPMW exhibits a sybtle asymmetry determined by the fractional difference in the values of thresholds. This prediction is testable by fitting such a curve to behavioral data in which agents are known to have some bias in their decision process that manifests as asymmetric decision thresholds or a non-zero starting point for their decison variable. The effects of drift rate, absolute threshold value, and threshold asymmetry are visually represented in Figure 3.

**Figure 3:**
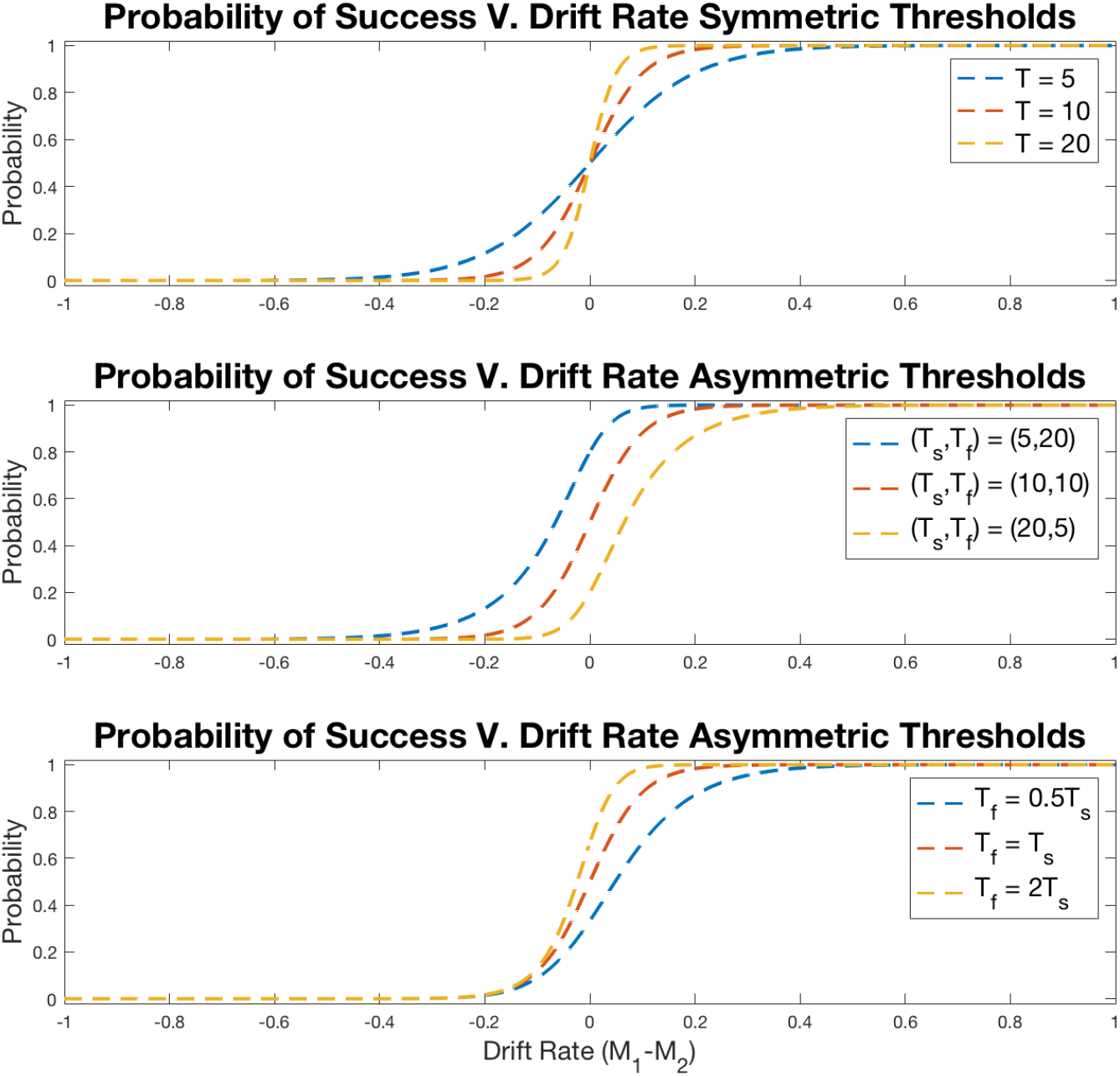
Probability of success versus drift rate curves under the free response paradigm. Subfigure 1 shows the effects of absolute value of thresholds on gain. Subfigure 2 shows the effects of relative absolute value of thresholds on bias. Subfigure 3 shows the effects of fractional differences in thresholds on asymmetry.

##### Mean Response Time

A useful parameter for describing an agent’s performance is the mean response time for both correct and incorrect responses, as opposed to the expected response time for each of them separately, which we will treat in the following section.

Finding the theoretical mean response time is done by setting absorbing states at the boundaries *T_f_* and *T_s_*. Next we find the characteristic matrix *N* = (*I* − *Q*)^*−*1^ where *Q* ∈ *R*^(*l−*2)*×*(*l−*2)^ s.t.

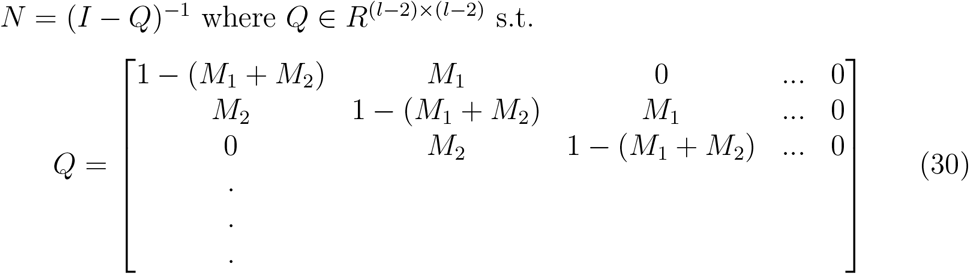

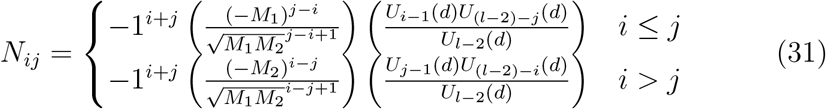

where *U_i_*(*d*) is the *i^th^* Chebyshev polynomial of the second kind, evaluated at *d*, and *d* is given by

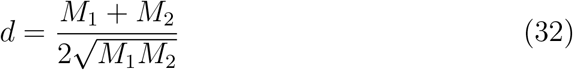

([9]). The sum over the entries of the *i^th^* row of *N* gives the expected time before absorption by either state, having started in state *i*. In the formulation we have developed, the agent always begins at *x*_0_ = 0, as asymmetries in the distances from starting points to thresholds are treated by the generalized definition of the thresholds. Therefore, we need only consider *i* = *T_f_*. Accordingly, the mean response time for the system is given by

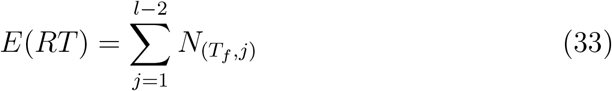

The terms of *N_ij_* depend on *M*_1_ and *M*_2_, but are also sensitive to their indices. This sensitivity is what makes the overall mean response time sensitive to the thresholds.

##### RT Distributions

The RT distributions for the free response paradigm are given by their first passage time across either threshold. The probabilities that the first passage time from initial state *S*_0_ to final states *s* and *f* (the absorbing states associated with the decision thresholds) is *t*, that we denote by *f_s_*(*t*) and *f_f_* (*t*) respectively. To find *f_s_*(*t*), we begin with the single-step transition probability matrix defined by *A* in eq. 17. Probability of absorption by success and failure thresholds respectively at time is given by

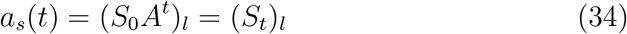

and

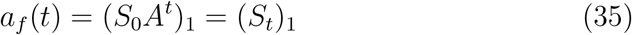

However, *a_s_*(*t*) and *a_f_* (*t*) give the probabilities of having been absorbed at time *t* or any preceding time. Therefore,

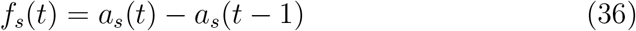

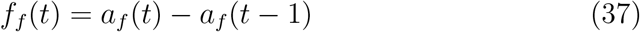

The tridiagonal nature of *A* allows closed forms of the distributions *f_s_*(*t*) and *f_f_* (*t*). To find these, we reduce *A* to matrix of matrices,

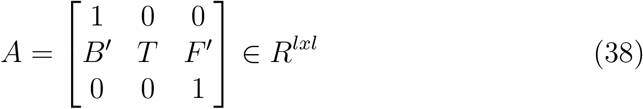

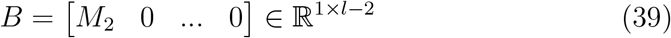

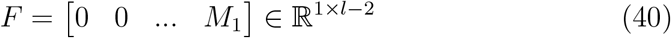

where *T* ∈ *R^l−^*^2*xl−*2^ is the familiar Toeplitz matrix used in the interrogation paradigm. Now,

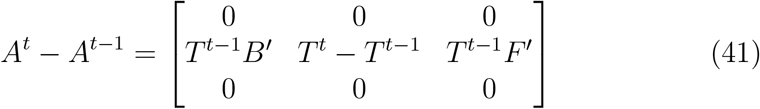

Because we are only interested in the probabilities of being in one of the two absorbing states, we are interested only in the top and bottom rows of this matrix, the *T_f_* + 1^*st*^ entries of which give the probability of succeeding or failing at time *t*:

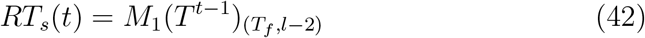

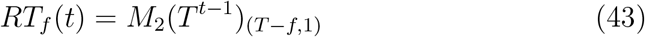

where the entries of *T^t−^*^1^ are given by eq. (13).

These distributions have inverse gaussian envelopes and exhibit period 2 oscillations, the peak amplitudes of which are dependent on the absolute value of *M*_1_ and *M*_2_ as well as their difference, *M*. As the upper and lower bounds of the gaussian envelope begin and end at zero, their differences, and therefore the amplitude of oscillations, are non-monotonic. By visual inspection, we find that the maximum oscillation amplitudes occur at the peak of the RT distributions. Three examples of RT distributions with different oscillation amplitudes are shown in figure 4.

**Figure 4:**
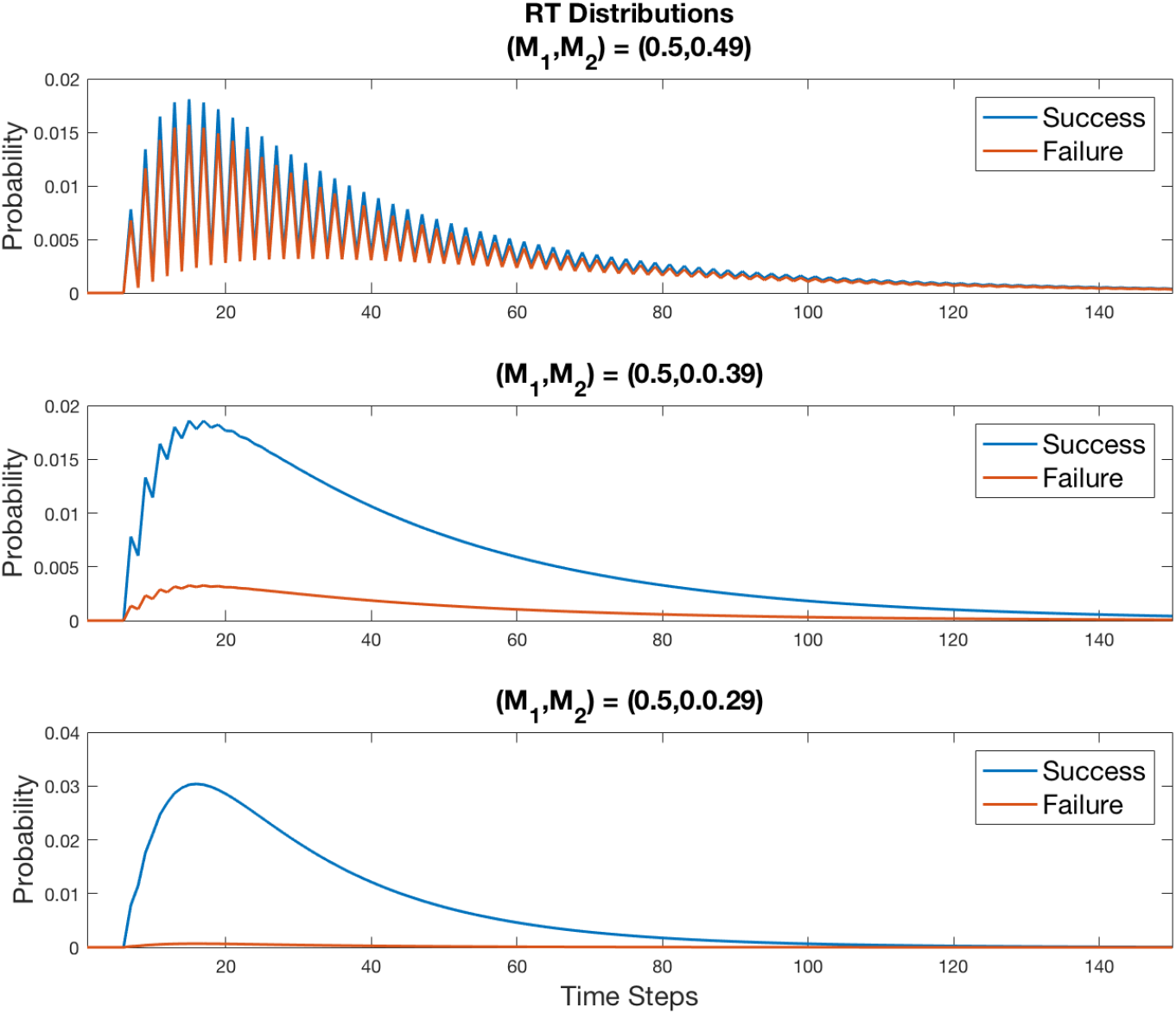
Example RT distribution curves. *M*_1_ remains constant across all three curves while *M*_2_ changes. The difference and the sum of *M*_1_ and *M*_2_ determine the amplitude of oscillations.

**Figure 5:**
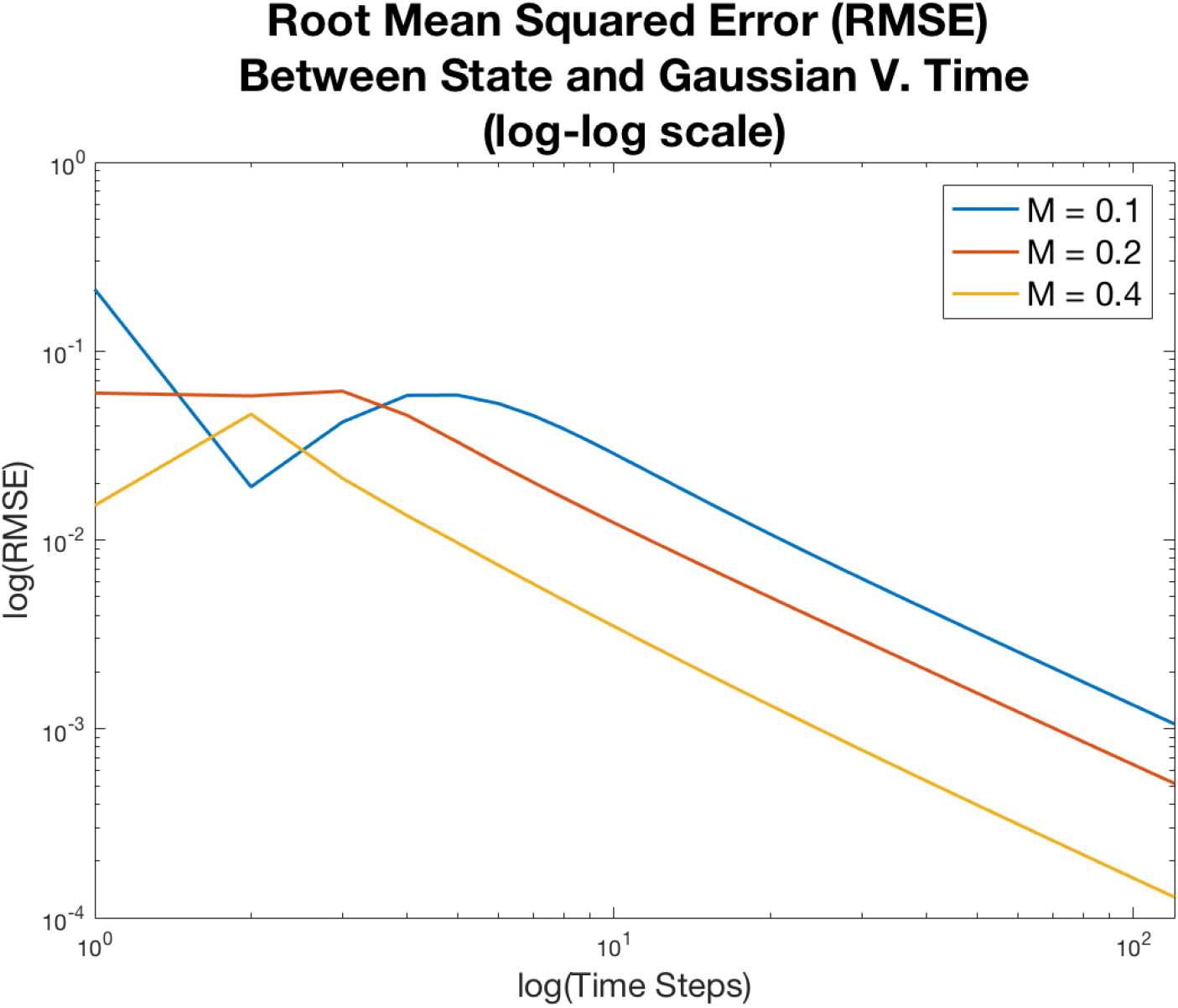
The root mean squared error (difference) between the state of a system and the gaussian distribution to which it converges. Note this plot is on a log-log scale.

We conduct a more thorough analysis of the oscillations in RT distributions in section 5.1.

#### 3.1.4 Convergence of the Single Integrator Model to a Traditional DDM

As we have established, the state of the system is given by a sum of IID random variables and, by the central limit theorem, we can assert that the success rate distribution of batches of free response trials, which is binomial at smaller numbers of batches, will converge to a gaussian such that

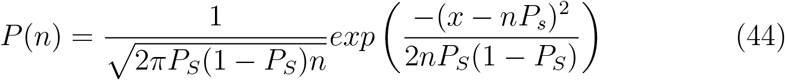

where *n* is the number of successes, and the probability of failure is taken to be 1 − *P_S_*, as non-responses are counted as failures.

Similarly, the probability distribution of the states for the interrogation paradigm (no absorbing states) as interrogation time increases is bounded by a gaussian distribution such that

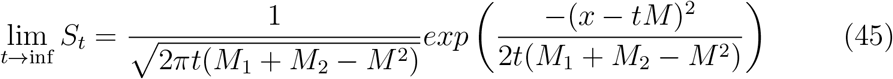

Thus, in the limit of sufficiently high thresholds, sufficiently long time, or (as we show in later sections) a sufficiently large number of integrators, the RT distributions are given by an inverse gaussian distribution describing the first passage times for the limiting state, a gaussian with the same drift rate *M* and variance (*M*_1_ + *M*_2_ − *M*^2^) as the system,

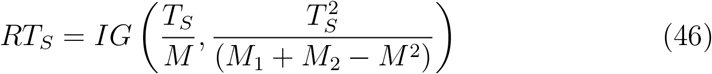

This system is identical to the traditional DDM, where the drift rate is given by

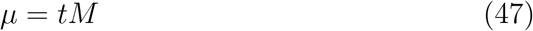

and variance given by

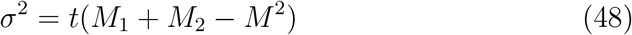

However, where the variance of the traditional DDM is a free parameter, in the MPMW model, the variance (denoted *σ*^2^ for brevity) is constrained to be a function of *M*_1_ and *M*_2_, and depends both on their relative value *M* and on their magnitude *M*_1_ + *M*_2_.

The variance of a system with a fixed drift rate *M* is bounded by a pair of values defined by *M*. We can find these by setting

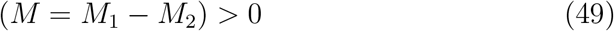

so

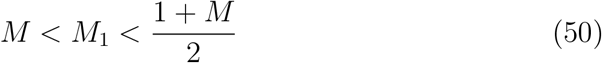

Therefore,

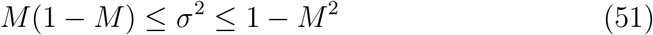

Thus, the MPMW distinguishes itself in an experimental context, as the relationship between the drift rate, variance, and MIEs for each well will be useful in fitting performance to generative landscapes to empirical data.

### 3.2 The Multi-Integrator Model

In the above treatment, we considered only the case of a single integrator used to make a categorical decision about the stimulus. However, it is reasonable to assume that a system with the capability of implementing multiple integrators (such as the brain) may benefit by doing so for signal averaging or summation, exploiting ergodicity by employing multiple parallel integrators for the same task. In the multi-integrator MPMW model, we consider such a system as a more biologically plausible extension to the MPMW single integrator model.

To do so, we assume a population of *m* integrators tasked with categorizing the stimulus based on its competing features. Let the set of MIEs for the stimulus components be given by *M*_1_, *M*_2_, representing the target (correct) and distractor (incorrect) stimuli respectively. We treat the system of integrators as a single sum integrator, the state of which, given by 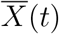, represents the total evidence accumulated for each alternative by each of the *m* integrators. The system may take a step of any integer size between [−*m, m*], with boundaries given by [−*T_f_, T_s_*]. This system is mathematically equivalent to a system that takes the average of accumulated evidence across all contributing integrators, the step size of which is bounded between [−1, 1], and may take on values that are integer multiples of 1/*m*, with boundaries [−*T_f_ /m, T_s_/m*]. We chose to represent the system by a sum integrator for notational clarity, but it remains conceptually identical. The state of the system is

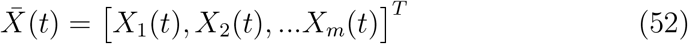

and evidence accumulates at each time step such that

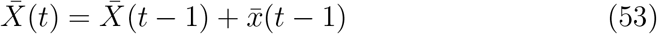

where

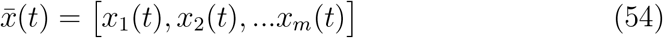

and each component *x_i_*(*t*) ∈ {−1, 0, 1}.

Let Δ_*m*_(*t*) denote the total evidence accumulated by the system of *m* integrators at time *t*. This is given by

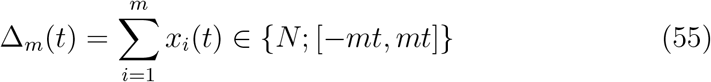

and the change in the total evidence at time *t* is given by a random variable

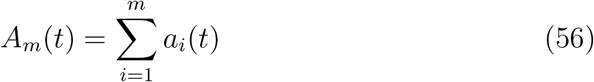

A decision is reached when the state of the system crosses either of two thresholds, (−*T_f_, T_s_*).

Let the number of integrators that have integrated evidence for the correct, incorrect, or neither option at a time step be given by *n_c_*, *n_i_*, and *n_o_* respectively.

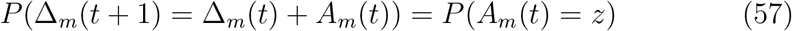

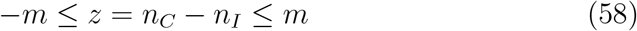

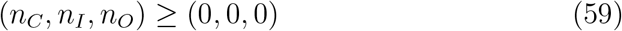

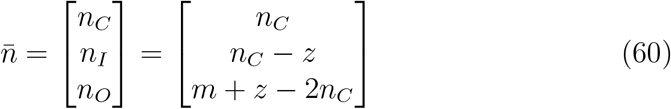

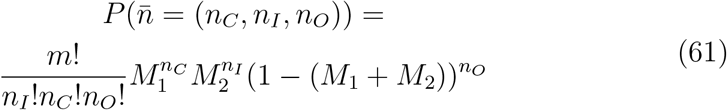

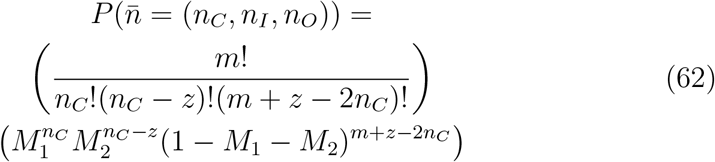

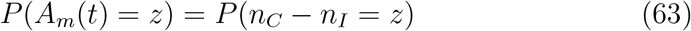

There can be more than one valid 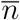 that yields a value *A* = *z*. The full transition probablility for a time step can be found algorithmically, but to do so requires the calculation of factorials, which become large quickly and therefore intractable to calculate. This can be avoided by calculating these values directly using the solutions for the Toeplitz tridiagonal matrices described in the interrogation paradigm section of the single integrator version of the model (section 3.1.1). It is straightforward to see that

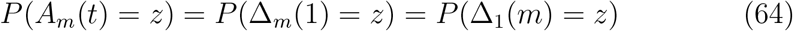

using the single step transition matrix for a single integrator, *T*, given by equation 6, such that

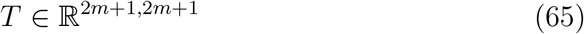

We find that the single step transition probabilities for a system of *m* integrators are given by

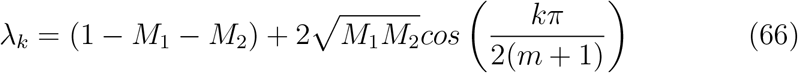

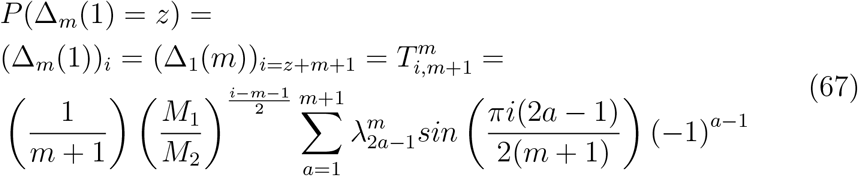

For notational clarity, these single step transition probabilities will be denoted by

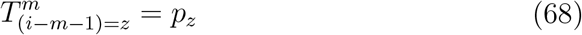

where

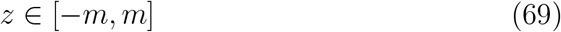

so that the probability that the state of the system after a single time step is

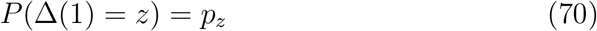

#### 3.2.1 The Interrogation Paradigm

For the multi-integrator system, the probability of success at a certain interrogation time can be found using the same method as described for the single integrator system in section 3.1.2. It is trivial to see that, where Δ_*m*_(*t*) is the the state of a system with number of integrators *m* at time *t*,

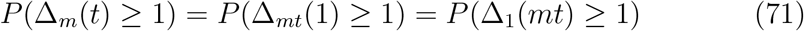

where Δ_*mt*_(1) is the probability of success *P_S_* under the interrogation paradigm for a single integrator system at time *mt*, and is given by

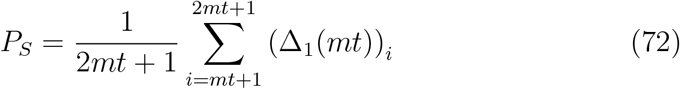

#### 3.2.2 The Free Response Paradigm

As in the case of the single integrator system, the free response paradigm for the multi-integrator system requires that we find the RT distributions for correct and incorrect responses at a given time. These are given by the probability distributions for the first crossing times at each threshold. Again, the thresholds are defined by absorbing states in the Markov chain. However, because the multiple integrator system may take up to *m* steps at a time, the single step transition matrix with absorbing states has a slightly more complex form. We may generate the single step transition matrix by directly computing the values of *p_z_* using the matrix *T* for a single integrator system at time *m*. Then, we can construct the single step matrix with absorbing states (analogous to eq 17), 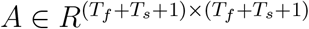.

The entries of the matrix *A* are as follows:

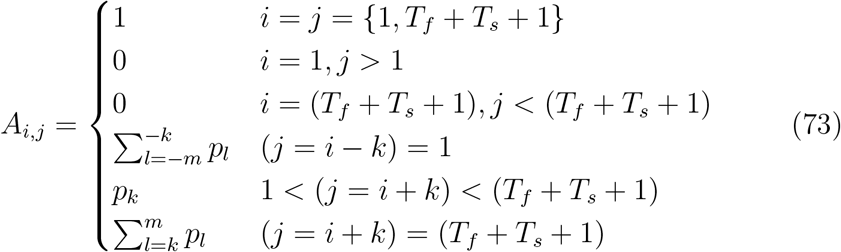

As before, the initial state of the system is Δ_*m*_(0) = 0, represented by 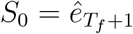. The state of the system at time *t* is *S_t_* = *S*_0_*T^t^*. The probability of absorption occurring at or before time *t* is 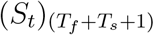 and the probability of the correct decision being made at time *t* is the probability of first crossing of the success threshold occurring at time *t*,

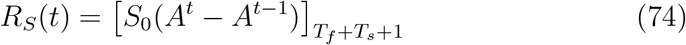

The probability of an incorrect decision at time *t* is the probability of first crossing of the failure threshold occurring at time *t*

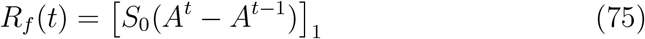

Unlike the single-integrator version of *A*, the multi-integrator transition matrix does not have a known closed-form solution of its powers, so the RT distributions must be calculated numerically. Probabilities of success and failure can be calculated by setting a time limit after which the trial ends and, if the agent has not responded, the trial is considered failed. In this case, the probability of success is given by

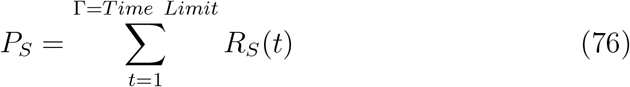

The total probability of failure is given by the probability of either responding incorrectly within the time limit or not responding at all. Probability giving an incorrect response is given by

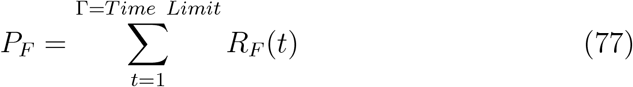

and

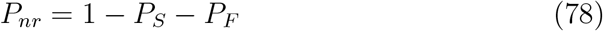

is the probability of failing to respond at all.

Finally, the mean response times for correct and incorrect responses are given by the inner product of the response time probability vectors over all time (*R_x_*(*t*)) and the associated time.

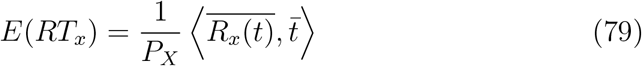

and the mean response time or mean time to absorption for either may be found either by calculating *N* = (*I* − *Q*)^*−*1^ or by taking

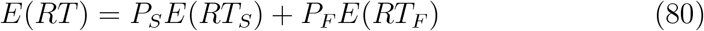

#### 3.2.3 Convergence of the Multi-Integrator Model to a Traditional DDM

In section 3.1.4, we showed that the single integrator model converges to a traditional DDM in the limit of long time, which occurs either with very high decision thresholds, very long decision times, or very high interrogation times. As we have discussed, in the multi-integrator model, number of integrators maps directly to the number of time steps in a single integrator model. Therefore, the multi-integrator MPMW model converges to a traditional DDM in the limit of large population. This property makes the MPMW framework one suitable for capturing similar qualitative properties of human behavior in perceptual decision making while offering a framework in which noise is an emergent property of the system and its context and drift rate is finite.

The PDF of the state of the decision variable in a single time step, in the limit of large integrator number of integrators is simply given by

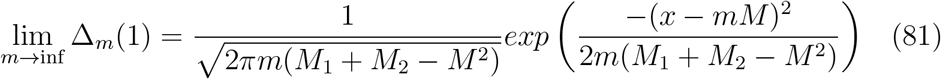

Also, in the limit of sufficiently high populations, the RT distributions are given by an inverse gaussian distribution for a gaussian with the same drift rate *mM* and standard deviation 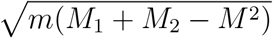 as the system,

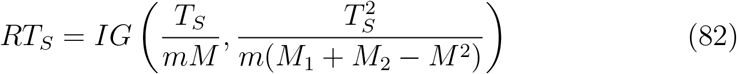

Just as in the case of a single integrator, this is identical to a classical DDM with drift rate given by

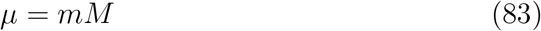

and variance given by

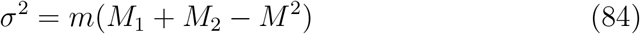

and the variance is takes on a range of values,

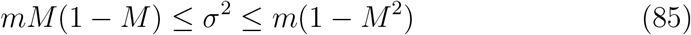

## 4 Numerical Results for the Multi-Integrator Interrogation Paradigm

### 4.1 Oscillations in the Probability of Success

As in the case of a single integrator, probability of success generally increases as a function of interrogation time, as long as *M*_1_ > *M*_2_; but it can exhibit both local and global non-monotonicities. For odd number of integrators, *P_S_* oscillates with a period of 2 time steps, which are bound within a collapsing envelope. To see why, we return to the equality

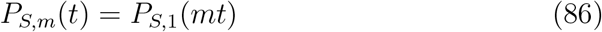

This equality states that the probability of success for a population of size *m* at each time *t* is the probability of success for a number of integrators 1 at every *m^th^* time step. For even populations, the sampling period, *m*, is an integer multiple of the oscillation period 2. Therefore, all samples will fall along the lower bound of the envelope for number of integrators 1. For odd populations, the sampling period, *m*, has value 2*k* + 1 where *k* is a positive integer value. As a result, samples fall along both upper and lower envelope bounds, with period 2, resulting in oscillations of period 2, with an amplitude that decreases more rapidly as number of integrators increases.

In an experimental setting, it is unlikely that it will be possible to directly map response time distributions to a single number of integrators. For example, an agent might recruit different numbers of integrators depending on circumstances across trials, blocks of trials, or experimental sessions. However, averaging across agents and time, we may assume that 50% of trials are performed with an odd number of integrators and 50% with an even number. In this case, period 2 oscillations may be visible but damped, as the measured probability of success curves would effectively be a linear combination of multiple theoretical probability of success curves across a spectrum of even and odd number of integrators.

#### 4.1.1 Numerical Analysis of Envelope Behavior as Number of Integrators Increases

The qualitative analysis outlined above can provide an expectation of the behavior of the envelope enclosing the oscillations in the probability of success function for the interrogation paradigm. Here, we explore this more rigorously.

For a fixed *M*, the upper and lower bounds of the envelope converge to a fixed value for all *m*. This is easy to see by taking the limit of

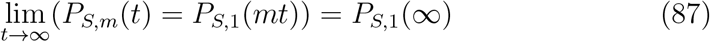

This equality also guarantees that the bounds of the envelope converge more rapidly as *m* increases, with the difference given by *P_S,_*_1_(*mt*) − *P_S,_*_1_(*mt* + *m*). Although the envelope bounds and differences can be found by direct computation, the equations are difficult to interpret. In order to more intuitively characterize envelope collapse, we attempted to fit them using two simple functional forms:

Fit 1, exponential: 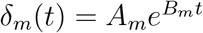

Fit 2, power law: 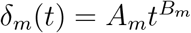

To perform the fits, we first determined if oscillations were present under the given conditions. We defined the oscillations of interest as having period 2 and therefore assumed that oscillations were present only if

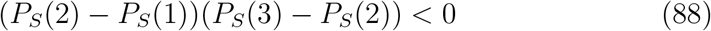

Under this circumstance, we characterized the envelopes by their difference as a function of time by assigning a dummy time, *t’* ∈ (*N*^+^ > 2). As the lower bound is only defined for odd times, we let

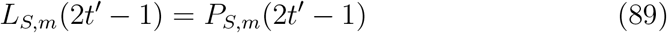

and approximate the lower bound at even times by

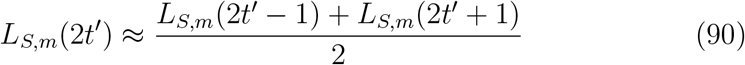

Similarly, the upper bound of the envelope is only defined for even times.

So we let

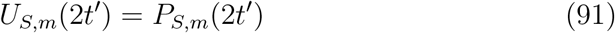

and approximate

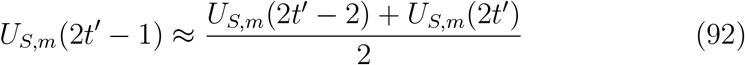

This leaves *U_S,m_*(*t’* = 1) not defined because it depends on *U_S,m_*(0). So we must begin with *t’* = 3. This allows us to define

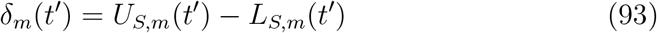

Both collapse to zero when *B_m_* < 0 as *t* → ∞.

Across five drift rates linearly distributed over [0.001, 0.1], and a range of 10 odd numbers of integrators [1, 19] ∈ *N_odd_*, we found that 56% of *P_S_*(*t*) functions exhibited oscillations. We then used nonlinear least squares regression to fit the difference between upper and lower bounds for those *P_S_*(*t*) that had oscillations. Defining the percent difference in adjusted *R*^2^ goodness of fit as

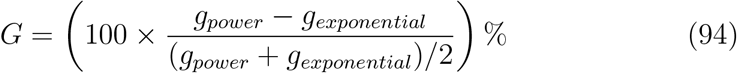

we found that 25% of curves were better fit by power laws than exponential, and that the mean 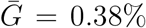 *G* and *median*(*G*) = −0.84%. Because oscillations diminished so quickly as a function of number of integrators, it was not possible to find a fit that best described *A_m_* and *B_m_* within these functions.

In order to determine whether oscillations might be measurable in human participants, we determined the combinations of drift rates and number of integrators required to observe oscillations. Specifically, we evaluated whether oscillations were observed and, if so, their maximum amplitude, for a set of 200 drift rates linearly distributed between *M* ∈ [0.001, 0.2], and 100 number of integrators ranging from *m* ∈ [1, 3, 5, *…*199]. For each drift rate, we found the largest number of integrators at which oscillations were identified, and fit the curve of that population against the drift rates using nonlinear least squares.

We found that the curve was best fit by a power law,

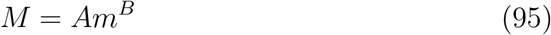

where *A* = 0.3884±0.0102, *B* = −0.9767±0.0133, with an adjusted *R*^2^ = 0.9957. According to this fit, at number of integrators exceeding 8, oscillations are present only at *M* < 0.05. This suggests that, at a biologically realistic number of integrators, the oscillations characteristic of this model will not be measurable using the interrogation paradigm.

In summary, we found that, in multi-integrator systems, the probability of success as a function of interrogation time exhibits period 2 oscillations when the number of integrators is odd. These oscillations are bound within an envelope that decays in time. The collapse of the envelope may be described either by a power law or an exponential. The largest number of integrators at which oscillations are identifiable is a power law of drift rate, and the fit of the power law suggests that, if a single neuron can be treated as an integrator, then at biologically plausible number of integrators, in the interrogation paradigm oscillations would likely not be observable, or would be heavily damped.

## 5 Free Response Paradigm Numerical Results

### 5.1 Oscillations in Response Time Distributions

As with the probabilities of success and failure in the interrogation paradigm, oscillations are present in the response times of the free response paradigm at a variety of drift rates and number of integrators. As described in section 3.1.3, response times are characterized by oscillations bound within an inverse gaussian envelope. By empirical observations, we found that, as with the interrogation paradigm, oscillations in the RT distribution have period 2 and are only present in odd-sized populations.

Also by empirical observation, and as can be seen in figure 4, we noted that the differences between the upper and lower envelopes were greatest at the peak of the distribution, the point at which the mean of the upper and lower bounds of the RT distribution’s inverse gaussian envelope has derivative = 0. Interestingly, this differs from the *P_S_*(*t*) distributions in the interrogation paradigm, where amplitude of oscillation decreases monotonically across the curve.

In order to determine whether oscillations were present in an RT distribution, we found the peak of the distribution and checked the derivative at surrounding points. At times preceding the peak, a true inverse gaussian curve would be monotonically increasing and have a strictly positive derivative. Similarly, at times after the peak, a true inverse gaussian would be monotonically decreasing and have a strictly negative derivative. We sought to identify oscillations of period 2, as those are the types of oscillations present in the single integrator model. To do so, we took the differences across five points, centered around the peak. For those both before and after the peak, we looked for differences of the pattern [*diff*_1_ < 0, *diff*_2_ > 0], and classified the curve as exhibiting oscillations if that pattern appeared either before or after the peak.

As with the interrogation paradigm, we sought to determine the maximum drift rate at which oscillations appeared for a range of number of integrators. We evaluated 300 drift rates between *M* = [5 × 10^*−*5^, 0.12]. We began by focusing on odd number of integrators and identified the largest drift rate at which oscillations were present (termed the “terminal drift rate”) either before or after the peak.

For 86.4% of the 22 number of integrators in which we found oscillations at the lowest drift rate, *M* ≥ 5 × 10^*−*5^, oscillations were present in the points following the peak at higher drift rates than for those preceding the peak. For populations at which oscillations were not present at drift rates *M* ≥ 5 × 10^*−*5^, the terminal drift rate value was simply set to *NaN*.

Unlike the interrogation paradigm, the curve mapping the terminal drift rates to their associated number of integrators was not monotonic. Rather, while it was generally decreasing, it exhibited several small peaks. To assess the general shape of the curve, ignoring its non-monotonicities, we used a weighted nonlinear least squares to fit the 22 populations that exhibited oscillations within the given range by setting weights to give high priority to the first and last points (*w* = 1), give low priority to points along an increasing portion of the curve associated with peaks (*w* = 0.25), and give medium priority to remaining points (*w* = 0.5). By setting these weight values, we were able to generate a fit that prioritized the monotonic portion of the curve, in an attempt to reduce the effects of the peaks. The resulting fit was a decreasing exponential given by

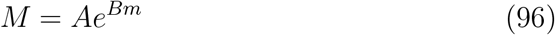

where *A* = 0.1164±0.0095 and *B* = −0.1823±0.0262, yielding an adjusted *R*^2^ = 0.9768.

This fit suggests that, at *M* < 0.05, oscillations in response time distributions will be detectable only at populations smaller than 5. The curve is similar in shape to the analogous curve for the presence oscillations in probability of success for the interrogation paradigm, as they arise from the same phenomenon of non-monotonicities in the PDF of the decision-state variable. In both cases, we can reasonably assume that, if a single integrator may represent a single neuron, then at biologically plausible number of integrators, oscillations would not be measurable.

In conclusion, our study of the oscillations predicted by the MPMW model suggests that, although the oscillations are fundamental to the model, if we assume that an integrator is mappable to a single neuron, oscillations are unlikely to be measurable in an experimental setting, as the brain uses very large populations of neurons, and oscillations cease to be measurable at number of integrators less than ten. This effect is similar to a scale effect upon the oscillations in preference predicted by the quantum random walk model of 2AFC ([13]). While the quantum random walk 2AFC model required a classical extension to account for these effects, they arise naturally within the MPMW framework. Other important components of the MPMW, such as noise, leak, and competition (mutual inhibition) among alternatives as emergent properties, its extensibility to an arbitrary number of alternatives, its coupling of parameters to automaticity and control, and its ability to differentiate among the effects thereof (⫾ eigenvalues paper), are independent of the result that oscillations diminish with scale.

## 6 Properties of and Performance at the Optimal Threshold

Among the advantages of the DDM is that it allows closed form analysis of thresholds that maximize reward rates given a set of trial conditions, thus allowing for normative predictions of behavior. The MPMW distinguishes itself from other quantum models in that, like the DDM, it too allows for optimal threshold analysis and captures similar effects to those found in [1] for the pure DDM. However because the MPMW framework’s definition of drift rate guarantees that it is finite as long as number of integrators is also finite, and because variance is a function of drift rate and MIEs rather than a free parameter, the MPMW exhibits some differences from the DDM in its properties surrounding optimal threshold. In this section, we consider analyses of the optimal threshold in the MPMW as a function of drift rate (where optimal threshold is defined as the threshold that maximizes reward rate, as in [1]), and compare its behavior with that observed for the DDM.

As in [1], we define the reward rate simply as

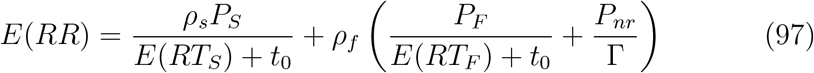

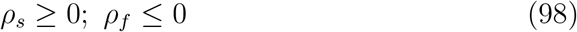

where *t*_0_ is the non-decision time, *ρ_i_* is the reward or penalty received for either success or failure on that trial, and Γ is the time limit. We assume that non-response trials receive the same penalty as failure trials, and that rewards and penalties remain static across trials.

One feature that distinguishes the MPMW from the DDM is the minimal non-decision time for each. In general, the minimum time associated with receiving a single bit of information and converting it to an action is labeled *t*_0_, and we assume it to be fixed. This is the non-decision time required for an agent to respond in a simple signal detection trial with very strong signal (i.e., 100% coherence). In the DDM, *t*_0_ accounts for the entirety of non-decision time. However, because the MPMW both discretizes time and places upper and lower bounds on drift rate (and therefore variance), an agent cannot respond before a time given by

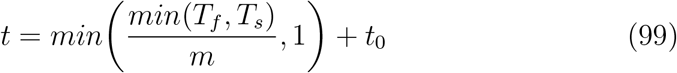

where *m* is the number of integrators applied to the task. This differentiates the MPMW from the standard DDM, or any other model in which drift rate is not bounded, in that it adds an additional parameter to the lower bound on decision time that scales with threshold. This may allow differentiation between DDM and MPMW models of decision-making: while the DDM can predict instantaneous decision times, the MPMW’s formalism imposes a threshold-dependent constant additive component of decision-time that is guaranteed regardless of signal strength or SNR.

An additional feature of the MPMW that distinguishes it from the DDM is that, under the MPMW, signal variance (”noise”) is inextricable from context. Variance is both a function of the difference between MIEs (drift rate), which refelcts conflict between the two stimuli, and their total magnitude, which reflects the probability of a dropped bit. Therefore, there is not a one-to-one mapping between drift rate and variance, but rather a range of variances that may be associated with each drift rate.This means that, when evaluating optimal threshold as a function of drift rate, variance as a function of drift rate must also be taken into account. That is, the fixed “relative variance” must be included in the evaluation, where relative variance is defined 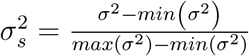. That is, relative variance describes where, in the spectrum of possible variances for a single drift rate, a specific variance falls. In figure 6, we have plotted variance versus drift rate for a range of five relative variances. Note that variance is a non-monotonic (quadratic) function of drift rate, and peak variance occurs at different drift rates for each relative variance, while minimum variance is always zero and always occurs at drift rate = 1. This means that, for different relative variances, maximum absolute variance has different values and occurs at different drift rates, but minimum absolute variance has the same value (0) and occurs at the same drift rate (1). We can intuit that a system will have minimal optimal threshold at minimal absolute variance, as high thresholds primarily serve to reduce error rate; and maximal optimal threshold at maximal absolute variance for the same reason. Based on this, we can postulate that a low relative variance will have its lowest optimal thresholds at low and high drift rate, with a peak optimal threshold occurring near its peak absolute variance, around *M* = 0.5. We can also anticipate that, as relative variance increases and the associated maximum variance both increases and occurs at lower drift rates, the peak optimal threshold will increase to reduce error rates and will occur at lower drift rates. Figure 6 shows that these intuitions are borne out by calculation.

**Figure 6:**
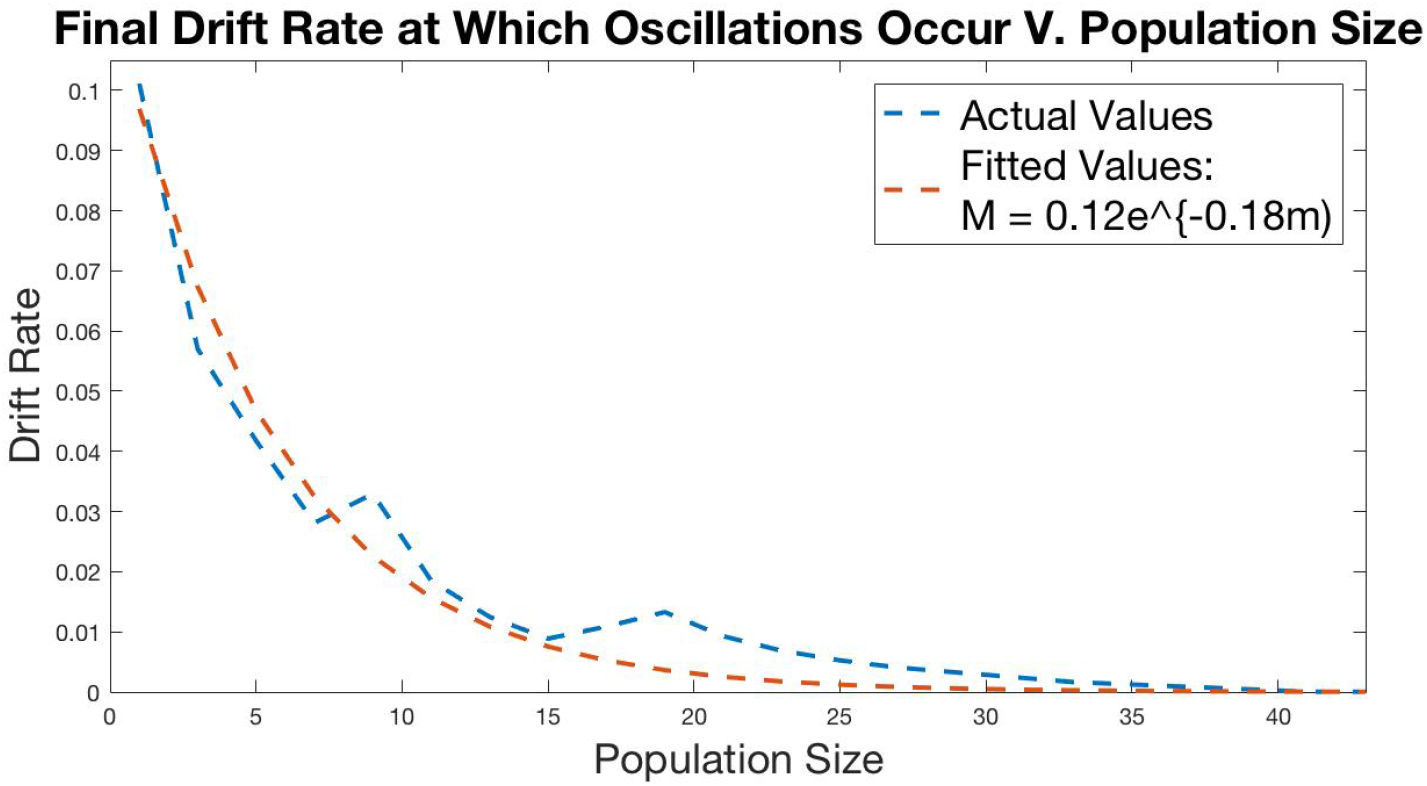
Maximum drift rates at which oscillations were measurably present in RT curves versus number of integrators. Note the presence of several peaks. The fit was designed to minimize the impact of these peaks on the overall fitted curve.

**Figure 7:**
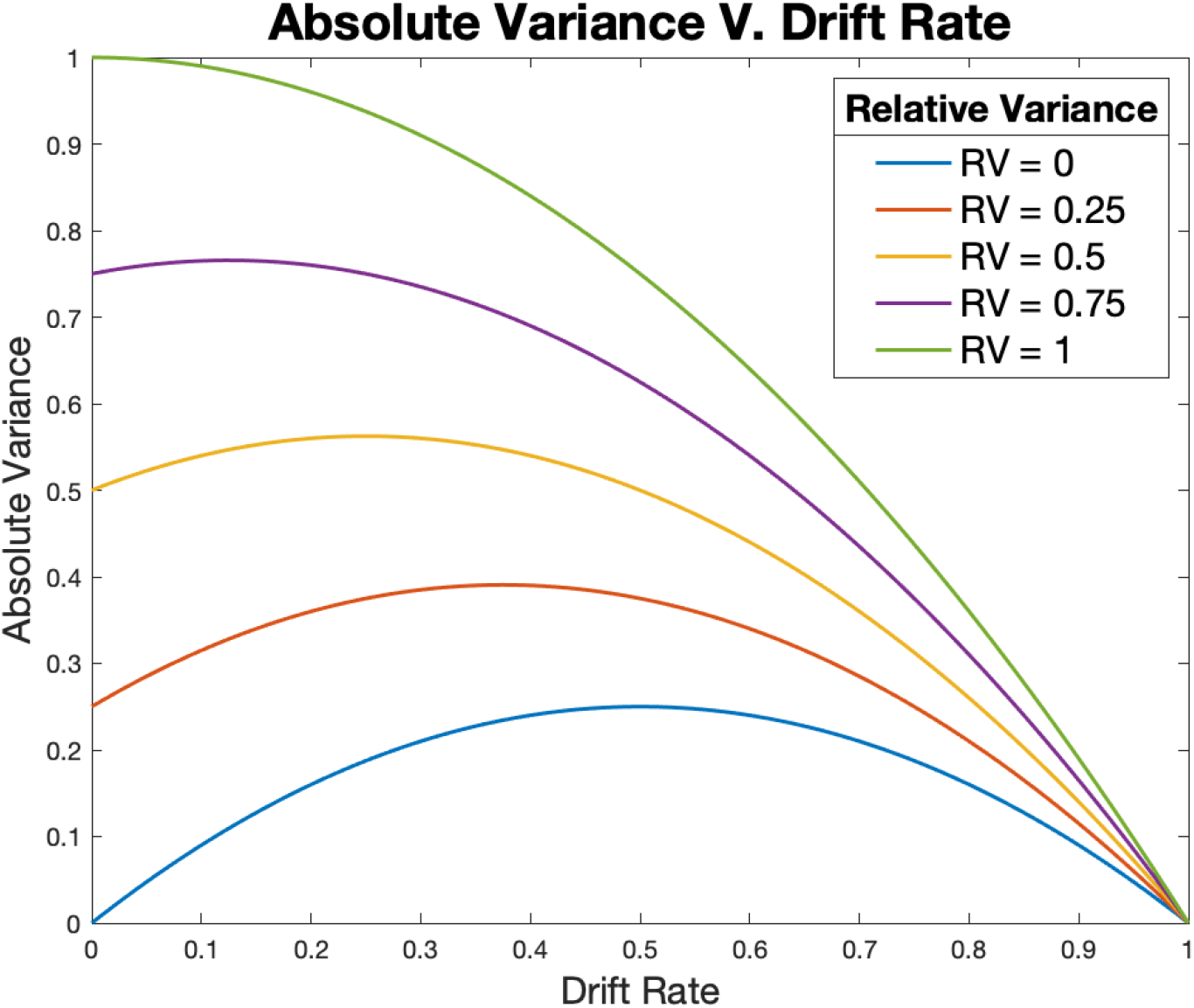
Absolute variance as a function of drift rate at multiple relative variances. Variance for a drift rate can occupy any value between relative variance = 0 and relative variance = 1.

**Figure 8:**
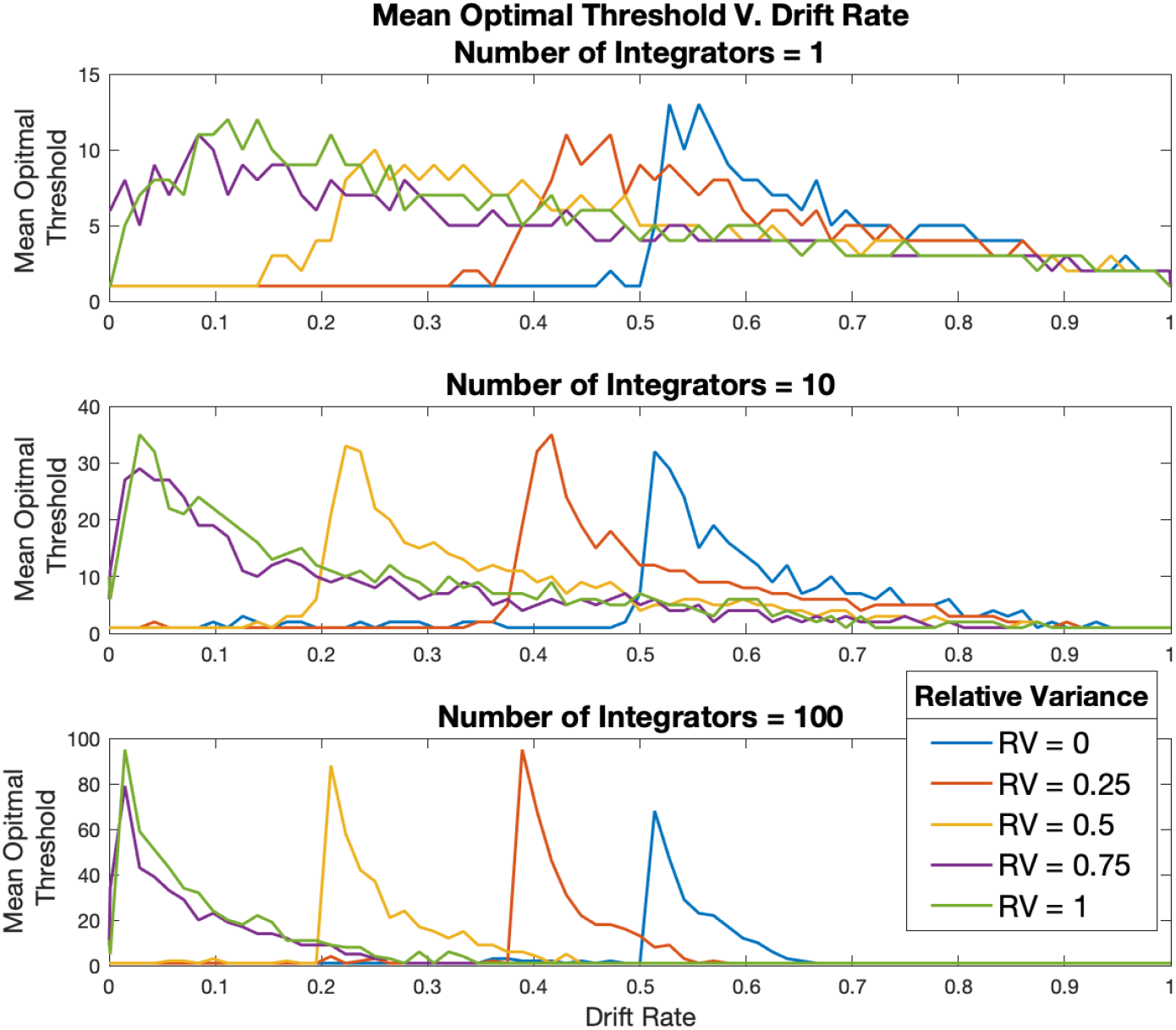
Mean optimal threshold as function of drift rate and relative variance for numbers of integrators 1, 10, & 100.

Although the MPMW allows direct theoretical computation of optimal threshold, performing that calculation on a computer is subject to limits on the computation of factorials in the case of direct calculation and floating point error in the case of matrix method calculations. In order to remove these challenges, we calculated the optimal threshold by simulating 20 batches of 100 trials of 2AFC tasks at each of 75 drift rates and 5 relative variances for thresholds ranging from 1 to 300 for population sizes 1, 10, 100. For each variance and drift rate, we defined optimal threshold to be the threshold that yielded the highest average reward rate for each batch, then formed the optimal threshold curve by taking the mean optimal threshold across the batches for all 5 variance sets and 75 drift rates.

As shown in figure 6, the mean optimal threshold as a function of drift rate curves followed a canonical shape similar to the one shown in [1]: minimal (approximately constantly one) at drift rates below those associated with maximum variance, rising steeply to a peak at maximum variance, and decaying gradually back to one. The peak was smaller and the decay from peak was more gradual for all curves at lower population numbers. Variance in the curves themselves was also higher at lower population numbers. Peaks were smaller and curves decayed more sharply for lower relative variance. All of these effects may be intuited by the fact that performance will be faster and, by properties of ergodicity, more consistent at higher population sizes, as well as at lower relative variance.

The curves are qualitatively similar to those of the DDM, but exhibited apparently sharper peaks and, because the MPMW places an upper bound on drift rate, did have an end point and could be shown to finitely, rather than asymptotically, decay to one. It is worth noting that the similarities between the curves presented in [1] and those produced by the MPMW are more visually clear at lower number of integrators than at higher numbers of integrators. This may be due to the fact that the curves produced by the MPMW are “zoomed out”, spanning the entire range of drift rates.

In conclusion, for the single integrator model, we have covered several differences between behavior of the DDM and the MPMW at optimal threshold, chief among which are the prediction of a precipitous decline in optimal threshold as a function of high drift rate, which emerges as a result of the finite value of drift rate in the MPMW, and the prediction of longer response times or low SNRs in the case of a weak signal, depsite a low optimal threshold. Both of these arise from the discrete treatment of time and the bounds on drift rate and SNR emerging from the mathematical formulation of *M*_1_ and *M*_2_ as probabilities. Because drift rate and variance for each drift rate is bounded in the MPMW but not in the DDM, and because the range of possible variances decreases as drift rate increases, the MPMW’s predictions may be testable by examining a range of signal strengths (SNR, in a dot motion task). In particular, experimenters may set SNR in such a task to some very high value and attempt to produce variance by some other means (not including the introduction of a distractor stimulus, as this would introduce a third well to the model and require a different analytical formulation). The MPMW predicts that data fits to such tasks should produce nearly zero variance. Similarly, fits to mid-range signal strengths should produce a larger range of variances than very high signal strengths and a smaller range of variances than very low signal strengths.

## 7 Discussion and Future Directions

In this work, we have presented the 2AFC application of the multi-particle multi-well (MPMW) quantum decision-making framework. The 2AFC model is largely tractable to closed-form analysis and mirrors key properties of classical DDMs, while offering a bridge between classical models and other quantum cognitive models pioneered by Busemeyer and colleagues (see [7]; [17]; [27]; [15]; [6]). In addition to capturing many qualitative properties of the DDM and convergence to a DDM within certain limits, the MPMW offers a powerful extension to the quantum cognitive framework aimed at modeling perceptual decision dynamics. There are several fundamental components of the MPMW framework that make it powerful in its intended capacity. First, by definition, the MPMW separates the quantum and classical components of the decision-making process (the state of the particles representing the incident stimulus and the state of the decision-variable), a property that makes it both analytically tractable and optimizable. Additionally, under the MPMW, variance (noise) is an emergent property of the full context of the decision-making process, including signal strength, the agent’s familiarity with a task and allocated control, as well as the agent’s state of arousal. Additionally, the MPMW incorporates several features of quantum systems not addressed by other quantum models, such as access of the quantum particles to the classically forbidden region, producing a property similar to “leak”, and mapping of energy eigenstates to an agent’s arousal state (treated in ⫾ eigenvalues paper). Also fundamental to the MPMW is the perception of stimuli presented in continuous time as discrete temporal packages, a model that has gained considerable support in recent years ([3]; [21]; [29]; [31]; [30]).

In this article, we showed that, in the limits of large time or a large number of integrators, the MPMW converges to a DDM. Outside of these limits the RT distributions of the MPMW model follow an inverse gaussian envelope, similar to the standard DDM, but exhibit period 2 oscillations arising secondarily from the quantum nature of the model, the competition between the integrators representing each alternative, and the probability that no information is integrated. Competition, which arises from the presence of finite depth attractor wells within the generative landscape, means that the decision variable’s PDF, after *t* = 0, is essentially broken into a tripartite set of “waves”, one at the starting point and one on either side of the starting point. Each wave has an amplitude that depends on *M*_1_ and *M*_2_. The wave to the right (positive *x*-axis) side of the starting point is associated with *M*_1_ and the particles integrated in favor of the target (correct) alternative. The wave to the left (negative *x*-axis) side of the starting point is associated with *M*_2_ and the particles integrated in favor of the distractor (incorrect) alternative. Finally, the wave centered at the starting point of the decision variable is associated with the probability that a particle is measured to be in the classically forbidden region and all associated “dropped bits”. As the system evolves, the component waves diffuse and interfere with each other.

Using the PDFs for the decision-variable, we described analytical solutions for probabilities of success and response times in both single and multi-integrator models, and showed that in the aforementioned limits of large scale, the state of the decision variable converges to a gaussian distribution, yielding RT distributions that are given by inverse gaussians comparable to those exhibited by the classical DDM, but with variance coupled to the effects of automaticity, salience, and control, rather than assumed as a free parameter. In short, the oscillatory effects that mark the MPMW in the single integrator model dissipate with large scale (in both time and number of integrators), just as quantum effects may be neglected in physical systems above a certain scale. Given that empirical data represents an aggregate of trials and individuals, it is unlikely that the oscillations of the MPMW model will be measurable. However, despite this, the MPMW provides a simple DDM-like model, the Markov properties of which make it easily extensible to larger numbers of alternatives and the quantum properties of which tie its parameters to such effects as automaticity, control, and arousal. Additionally, the boundedness of drift and dependence of SNR on *M*_1_ and *M*_2_ make distinguishable predictions beteween the behaviors of the DDM and MPMW at optimal threshold.

Although other quantum models also exhibit interference in the PDF of the decision variable, it has not been characterized as periodic in any specific frequency, but emerges as the components of the PDF for the single particle reflect at the boundaries of its well, which are also the decision boundaries. A distinguishing characteristic of the MPMW model is that it separates the quantum particles (which represent information from incident stimuli) from the classical decision variable and treats decision boundaries (thresholds) as absorbing rather than reflecting. Thus, in the interrogation paradigm, the decision variable is not subject to boundaries at all, and so it does not predict that an agent will change their mind over time if repeatedly questioned. Rather, in the MPMW, damped oscillations appear only in the probability of success at different interrogation times, a curve that assumes the agent will only be questioned once. As stated above, these oscillatory effects are subject to decay as the system’s time or population scale increases. A similar effect upon the predicted oscillations in preference is borne out by empirical research intended to test the quantum random walk model of 2AFC ([13]). While the quantum walk 2AFC model required a classical extension to account for these effects, they arise naturally within the MPMW framework.

The analyses of the MPMW presented here suggest several promising directions for further investigation. One, currently under pursuit, is the expansion of the two alternative double-well formulation to the multi-alternative multi-well formulation. This straightforward expansion uses the same methods of solving the TISE as are used in the 2AFC case and is minimally more complex. For *N* alternatives, the MPMW simply assigns each alternative to one of *N* finite square wells with width determined by stimulus salience or task automaticity and depth determined by allocated cognitive control. Thus, the MPMW can be used to include as many wells within the landscape as alternatives from which an agent must select. Each well produces (by solving the TISE) an associated probability that a bit of incident information will be integrated as evidence for that alternative. Evidence accumulation may be represented by a set of markov chains, and the differences between the states of each chain determine decision time and probability of success. Under fixed landscape parameters (as assumed in this article), this model is trivial to simulate, making analysis of its predictions relatively easy. This feature distinguishes the MPMW from other models of decision dynamics.

Another important feature of the MPMW is that it provides a natural interpretation of the effects of automaticity and control in terms of the width and depth of wells, respectively, as well as arousal in terms of energy eigenstates. This provides a novel approach to separating and analyzing the effects of practice and attention on performance ⫾ (eigenvals paper). For example, relaxing the assumption of static landscapes (well parameters) can incorporate the effects of reallocation of control both within and between trials, manifesting as changes in drift rate and variance of the decision process. This offers the possibility of an analytically tractable approach analogous to understanding the effects of variable drift rate ([19]; [20] [14]; [36]) and collapsing decision bounds ([26]; [10]) that have been addressed by extensions to the classical DDM, but proven a challenge to formal analysis. Among the effects of variable drift rates, we anticipate changes in the presence and amplitude of oscillations in the PDF of the decision variable, probability of success in the interrogation paradigm, and RT distributions. Finally, relaxing the assumption of equally probable bound states, provide a formally rigorous and analytically tractable approach to accounting for the effects of arousal on performance in terms of the eigenenergies of the system. Combining this with an analysis of how asymmetries in well depth and width effect eigenenergy spectra may provide a unifying, formally explicit account of interactions between automaticity, control and arousal — factors that are widely assumed to interact in affecting performance, but that are often treated in isolated or only qualitative form.

## 8 Conclusions

In conclusion, we have shown that the MPMW framework applied to 2AFC offers a quantum approach to perceptual decision-making that stands as a powerful alternative to the quantum random walk 2AFC [7] while coexisting with quantum cognitive models that have been used to describe a number of otherwise poorly understood effects. By its use of both quantum and classical variables, the MPMW marries the quantum cognitive framework with simpler classical versions of the DDM model using standard linear and nonlinear dynamical systems approaches. This approach allows stochasticity to arise as a fundamental emergent property of the system determined directly by the parameters of the generative landscape’s attractor wells, which reflect task parameters and the agent’s experience, rather than requiring noise to be introduced as an external variable. In the limits of long time or large number of integrators, the MPMW converges to a classical DDM with variance determined by system parameters rather than fitting. The oscillations it exhibits in RT distributions and probability of success under interrogation arise from interference in the state of the decision variable, a quantum effect that, much like quantum effects in physical systems, dissipates in large scale systems. We have also indicated ways in which, in future work, the MPMW model can be extended by considering the effects of landscape parameters on eigen state distributions, providing the potential for an integrated and formally rigorous approach to interactions between automaticity, control, and arousal on performance in N-alternative forced choice decision making tasks.

## References

[1] Rafal Bogacz et al. “The Physics of Optimal Decision Making: A Formal Analysis of Models of Performance in Two-Alternative Forced-Choice Tasks”. English. In: Psychological review 113.4 (2006), pp. 700–765.

[2] Kenneth H. Britten et al. “Responses of neurons in macaque MT to stochastic motion signals”. English. In: Visual neuroscience 10.6 (1993), pp. 1157–1169.

[3] Timothy J. Buschman and Earl K. Miller. “Shifting the spotlight of attention: evidence for discrete computations in cognition”. English. In: Frontiers in human neuroscience 4 (2010), pp. 194–194.

[4] J.R. Busemeyer and J.T. Townsend. “Decision field theory : a dynamic-cognitive approach to decision making in an uncertain environment”. English. In: Psychological review 100.3 (1993), pp. 432–459.

[5] Jerome R Busemeyer, Zheng Wang, and Ariane Lambert-Mogiliansky. “Empirical comparison of Markov and quantum models of decision making”. In: Journal of Mathematical Psychology 53.5 (2009), pp. 423–433.

[6] Jerome R. Busemeyer and Zheng Wang. “What Is Quantum Cognition, and How Is It Applied to Psychology?” English. In: Current directions in psychological science : a journal of the American Psychological Society 24.3 (2015), pp. 163–169.

[7] Jerome R. Busemeyer, Zheng Wang, and James T. Townsend. “Quantum dynamics of human decision-making”. English. In: Journal of mathematical psychology 50.3 (2006), pp. 220–241.

[8] V. Feller and W. Feller. An Introduction to Probability Theory and Its Applications, Volume 1. A Wiley publication in mathematical statistics. Wiley, 1968. isbn: 9780471257080. url: https://books.google.com/books?id=ZfFQAAAAMAAJ.

[9] C. M. d. Fonseca and J. Petronilho. “Explicit inverse of a tridiagonal k-Toeplitz matrix”. English. In: Numerische Mathematik 100.3 (2005), pp. 457–482.

[10] Peter Frazier and Angela J Yu. “Sequential Hypothesis Testing under Stochastic Deadlines”. In: Advances in Neural Information Processing Systems. Ed. by J. Platt et al. Vol. 20. Curran Associates, Inc., 2008. url: https://proceedings.neurips.cc/paper/2007/file/9c82c7143c102b71c593d98d96093fde-Paper.pdf.

[11] Henry Hamburger. Donald R. J. Laming. Information, theory of choice-reaction times. New York: Academic Press, 1968. English. 1969.

[12] Davod Khojasteh Salkuyeh. “Positive integer powers of the tridiagonal Toeplitz matrices”. In: International Mathematical Forum 1 (Jan. 2006).

[13] Peter D. Kvam, Jerome R. Busemeyer, and Timothy J. Pleskac. “Temporal oscillations in preference strength provide evidence for an open system model of constructed preference”. English. In: Scientific reports 11.1 (2021), pp. 8169–8169.

[14] Yuan S. Liu, Philip Holmes, and Jonathan D. Cohen. “A Neural Network Model of the Eriksen Task: Reduction, Analysis, and Data Fitting”. English. In: Neural computation 20.2 (2008), pp. 345–373.

[15] Andrzej Lukasik. “Quantum models of cognition and decision”. English. In: International journal of parallel, emergent and distributed systems 33.3 (2018), pp. 336–345.

[16] Darren Pais et al. “A mechanism for value-sensitive decision-making”. English. In: PloS one 8.9 (2013), e73216–e73216.

[17] Emmanuel M. Pothos and Jerome R. Busemeyer. “A quantum probability explanation for violations of ‘rational’ decision theory”. English. In: Proceedings of the Royal Society. B, Biological sciences 276.1665 (2009), pp. 2171–2178.

[18] Roger Ratcliff. “A theory of memory retrieval”. English. In: Psychological review 85.2 (1978), pp. 59–108.

[19] Roger Ratcliff and Jeffrey N. Rouder. “Modeling Response Times for Two-Choice Decisions”. English. In: Psychological science 9.5 (1998), pp. 347–356.

[20] Roger Ratcliff, Trisha Van Zandt, and Gail McKoon. “Connectionist and Diffusion Models of Reaction Time”. English. In: Psychological review 106.2 (1999), pp. 261–300.

[21] John H. Reynolds, Leonardo Chelazzi, and Robert Desimone. “Competitive Mechanisms Subserve Attention in Macaque Areas V2 and V4”. English. In: The Journal of neuroscience 19.5 (1999), pp. 1736–1753.

[22] Morgan Rosendahl, Anastasia Bizyaeva, and Jonathan Cohen. “A Novel Quantum Approach to the Dynamics of Decision Making”. In: (2020).

[23] Michael N. Shadlen and William T. Newsome. “Motion Perception: Seeing and Deciding”. English. In: Proceedings of the National Academy of Sciences - PNAS 93.2 (1996), pp. 628–633.

[24] Michael N. Shadlen and William T. Newsome. “The Variable Discharge of Cortical Neurons: Implications for Connectivity, Computation, and Information Coding”. English. In: The Journal of neuroscience 18.10 (1998), pp. 3870–3896.

[25] Mervyn Stone. “Models for choice-reaction time”. English. In: Psychometrika 25.3 (1960), pp. 251–260.

[26] Satohiro Tajima, Jan Drugowitsch, and Alexandre Pouget. “Optimal policy for value-based decision-making”. English. In: Nature communications 7.1 (2016), pp. 12400–12400.

[27] Jennifer S. Trueblood and Jerome R. Busemeyer. “A Quantum Probability Account of Order Effects in Inference”. English. In: Cognitive science 35.8 (2011), pp. 1518–1552.

[28] Marius Usher and James L. McClelland. “The Time Course of Perceptual Choice: The Leaky, Competing Accumulator Model”. English. In: Psychological review 108.3 (2001), pp. 550–592.

[29] Rufin VanRullen. “Perceptual Cycles”. English. In: Trends in cognitive sciences 20.10 (2016), pp. 723–735.

[30] Rufin VanRullen, Thomas Carlson, and Patrick Cavanagh. “The Blinking Spotlight of Attention”. English. In: Proceedings of the National Academy of Sciences - PNAS 104.49 (2007), pp. 19204–19209.

[31] Rufin VanRullen et al. “Attention-Driven Discrete Sampling of Motion Perception”. English. In: Proceedings of the National Academy of Sciences - PNAS 102.14 (2005), pp. 5291–5296.

[32] A. Wald. Sequential Analysis. Dover phoenix editions. Dover Publications, 2004. isbn: 9780486439129. url: https://books.google.com/books?id=oVYDHHzZtdIC.

[33] Xiao-Jing Wang. “Probabilistic Decision Making by Slow Reverberation in Cortical Circuits”. English. In: Neuron (Cambridge, Mass.) 36.5 (2002), pp. 955–968.

[34] Zheng Wang and Jerome R. Busemeyer. “A Quantum Question Order Model Supported by Empirical Tests of an A Priori and Precise Prediction”. English. In: Topics in cognitive science 5.4 (2013), pp. 689–710.

[35] Zheng Wang et al. “Context effects produced by question orders reveal quantum nature of human judgments”. English. In: Proceedings of the National Academy of Sciences - PNAS 111.26 (2014), pp. 9431–9436.

[36] Angela J. Yu, Peter Dayan, and Jonathan D. Cohen. “Dynamics of Attentional Selection Under Conflict: Toward a Rational Bayesian Account”. English. In: Journal of experimental psychology. Human perception and performance 35.3 (2009), pp. 700–717.

